# Medial Temporal Default Mode Network Selectively Encodes Autobiographical Visual Imagery

**DOI:** 10.1101/2025.11.25.690576

**Authors:** Andrew J. Anderson, Adam G. Turnbull, Feng V. Lin

## Abstract

The human brain’s capacity to imagine visual scenes from memory is thought to rely on the medial temporal subsystem of the default mode network (MT-DMN), yet the neural codes supporting this ability remain poorly understood. We combined functional magnetic resonance imaging (fMRI) with vision and language artificial intelligence models to characterize neural codes during autobiographical imagination. Fifty participants imagined reexperiencing twenty natural scenarios while undergoing fMRI, when cued by generic text prompts (e.g., wedding, exercising, driving). Individual scenes were modeled using Stable Diffusion to generate personalized synthetic images from verbal descriptions of the scenarios imagined, collected beforehand. These depictions were then transformed into image-recognition network embeddings. Representational Similarity Analysis revealed that the MT-DMN encoded the participant-specific representational structure of image embeddings, even when controlling for semantic features derived from a large language model. This effect was absent in other networks and during reading without imagination, identifying the MT-DMN as a core substrate for the visual reconstruction of autobiographical experiences.

## Introduction

Through imagination, the human brain reconstructs visual scenes from memory, synthesizing the appearance of places, people, and objects, even when these mental images are vague (Pearson et al., 2019). The medial temporal subsystem of the default mode network (MT-DMN; Raichle et al., 2019), encompassing parahippocampal, retrosplenial, and lateral parietal cortices, is thought to support such imagery because it engages during tasks involving memory, imagination, and spatial processing (Andrews-Hanna et al., 2010, 2014; Andrews-Hanna and Grilli, 2021). However, the neural codes that prospectively encode the reconstruction of visual scenes within MT-DMN remain unknown. Identifying these codes requires a representational model of the idiosyncratic scenes that individuals imagine, which is a challenge, further compounded by the need to disentangle visual from non-visual components such as affect, dialogue, and social relevance. Here, we address these challenges by combining a state-of-the-art text-to-image AI model (Stable Diffusion; Rombach et al., 2022; Podell et al., 2023) to model imagined scenes from verbal descriptions, with a large language model (Radford et al., 2019) to capture accompanying non-visual semantic content. This approach enables a characterization of the neural correlates of self-generated, participant-specific visual imagery, and an evaluation of their specificity to MT-DMN.

Despite extensive research into self-generated mental states and the DMN (Buckner and DiNicola, 2019; Smallwood et al., 2021; Yeshurun et al., 2021; Menon, 2023), methods for isolating neural representations that reflect internally generated visual scene simulations have been limited. Subjective mental imagery has typically been studied using experience sampling, in which participants report whether their ongoing thoughts are visual or verbal in form (Delamillieure et al., 2010; Gorgolewski et al., 2014; Smallwood and Schooler, 2015; Sormaz et al., 2018). However, such coarse self-reports cannot distinguish among the diverse visual scenes that individuals imagine. Likewise, although studies of episodic recall and prospection have successfully differentiated neural representations associated with distinct events (Chadwick et al., 2010; Baldassano et al., 2017; Chen et al., 2017; Anderson et al., 2020), their spatial or temporal context (Bonnici et al., 2012; Nielson et al., 2015), and the people or objects they involve (Thornton et al., 2017; Silson et al., 2019; Sreekumar et al., 2018; Benoit et al., 2019; Gagnepain et al., 2020; Gilmore et al., 2021), they have not identified neural correlates of the idiosyncratic visual details defining imagined scenes.

In contrast, neurocomputational accounts of mental imagery for single entities - such as animals, tools, people, and buildings - are comparatively well developed (Robinson et al., 2023; Pearson, 2019). Following early advances in modeling visual object perception (Leeds et al., 2013; Yamins et al., 2014; Khaligh-Razavi and Kriegeskorte, 2014; Horikawa and Kamitani, 2017; Nonaka et al., 2021) using computer vision systems (e.g., Lowe, 2004; Simonyan and Zisserman, 2015), similar approaches have been extended to model mental imagery of pre-viewed object images (Horikawa and Kamitani, 2017; Breedlove et al., 2020) and object nouns (Anderson et al., 2015; Soto et al., 2020). A general finding across imagery and perception is that activation patterns along the ventral visual pathway - central to object recognition (Ungerleider and Mishkin, 1982; Ungerleider and Haxby, 1994) - correlate with image embedding vectors extracted from computational object-recognition models (Khaligh-Razavi and Kriegeskorte, 2014; Eickenberg et al., 2017; Horikawa and Kamitani, 2017; Nonaka et al., 2021; Gong et al., 2023). These embeddings reflect high-level visual correlates of semantic categories, such as objects, animals, landmarks, and other entities, that abstract over low-level variation in position, viewpoint, and lighting. Notably, such embeddings also predict fMRI responses to viewing natural scenes within scene-selective regions (Groen et al., 2018) that overlap with the MT-DMN. These regions encode object categories (Stansbury et al., 2013; Çukur et al., 2016), contextual associations among scene elements (Bar, 2003, 2004; Aminoff et al., 2013), and background texture information (Cant and Xu, 2012). Indeed, the relative sensitivity of image recognition models to object context, and texture over shape (Geirhos et al., 2019; Jacob et al., 2021; Bowers et al., 2023) may even render them advantageous for modeling activity in scene-selective cortices.

Modeling visual imagery associated with autobiographical experiences presents a unique challenge because no objective ground truth for mental images exists. Although some episodic memories are photographically documented, what individuals recall or imagine often diverges from reality by blending elements from distinct events or introducing imagined details (Schacter et al., 2007a; Addis et al., 2007; Schacter et al., 2007b, 2012; Szpunar et al., 2007). Recent advances in text-to-image AI systems, such as Stable Diffusion (Rombach et al., 2022; Podell et al., 2023) now offer a means to approximate these internal visualizations by generating images directly from participants’ verbal descriptions. The present study leverages this approach to test whether activation patterns within the MT-DMN selectively encode participant-specific (autobiographical) visual structure, and to determine whether such structure is absent in other brain networks or when imagination is not engaged. In doing so, this work advances understanding of the neural basis of imagination and memory and establishes image-generation AI as a methodological complement to emerging language-model–based neural-decoding approaches (e.g. Nishida et al. 2018, Tang et al. 2023, Horikawa 2025, Matsuyama et al. 2025), capable of distinguishing internally generated visual imagery

## Results

To test the hypothesis that the MT-DMN selectively encodes participant-specific visual structure during autobiographical imagination, we reanalyzed a previously collected fMRI dataset from 50 adults (Anderson et al., 2020; Wang et al. 2024, **Fig, 1A**). The sample included 25 young adults (mean±SD age = 24±3 years; 16F, 9M) and 25 older adults (mean±SD age= 73±7 years; 16F, 9M). Although age-related differences were not a focus of the current study, analyses were conducted both across and within age groups to establish the generalizability of findings.

Prior to scanning, participants engaged in a standardized imagery task in which they vividly imagined 20 fixed, experimenter-selected naturalistic scenarios (e.g., exercising, shopping, wedding), drawing from their own autobiographical experiences. Each participant then provided a brief verbal description of their mental image (Anderson et al., 2020), which served as the input for later computational modeling.

To further characterize the content of these scenarios, and the extent to which they reflected episodic memories (Tulving et al. 1972, Wheeler et al. 1997), participants rated each on: (1) whether the event described had occurred in real life, and (2) the vividness of the imagined scene. These ratings suggested that most scenarios were based on real-life events (mean rating: 5.6 and 5.5 for young and elderly participants, respectively; scale: 0-6) and were vividly imagined (mean vividness: 4.6 and 5.2, respectively, scale 0-6).

Later, participants underwent fMRI, where they re-imagined the same 20 scenarios, presented in a randomized order, five times each. Scenarios were cued with generic written prompts that contained no participant-specific information (e.g., “A dancing scenario”). Standard fMRI preprocessing was conducted using fMRIPrep (Esteban et al., 2019), and scenarios repetitions were averaged to yield a single fMRI volume per scenario per participant.

### Modeling participant-specific autobiographical mental images

To model participant-specific scene visualizations, Stable Diffusion 2.1 (Rombach et al., 2022) was used to generate five synthetic images per scenario for each participant, based on their verbal descriptions of the autobiographical scenarios (Fig. 1A). The resulting images, available online as **Supporting Data**, generally appeared photorealistic, when viewed from a moderate distance, and typically featured coherent combinations of backgrounds, actors, and objects arranged in plausible spatial configurations. Fine details such as facial features or limb positions were often distorted, consistent with known limitations of Stable Diffusion.

**Figure 1.**
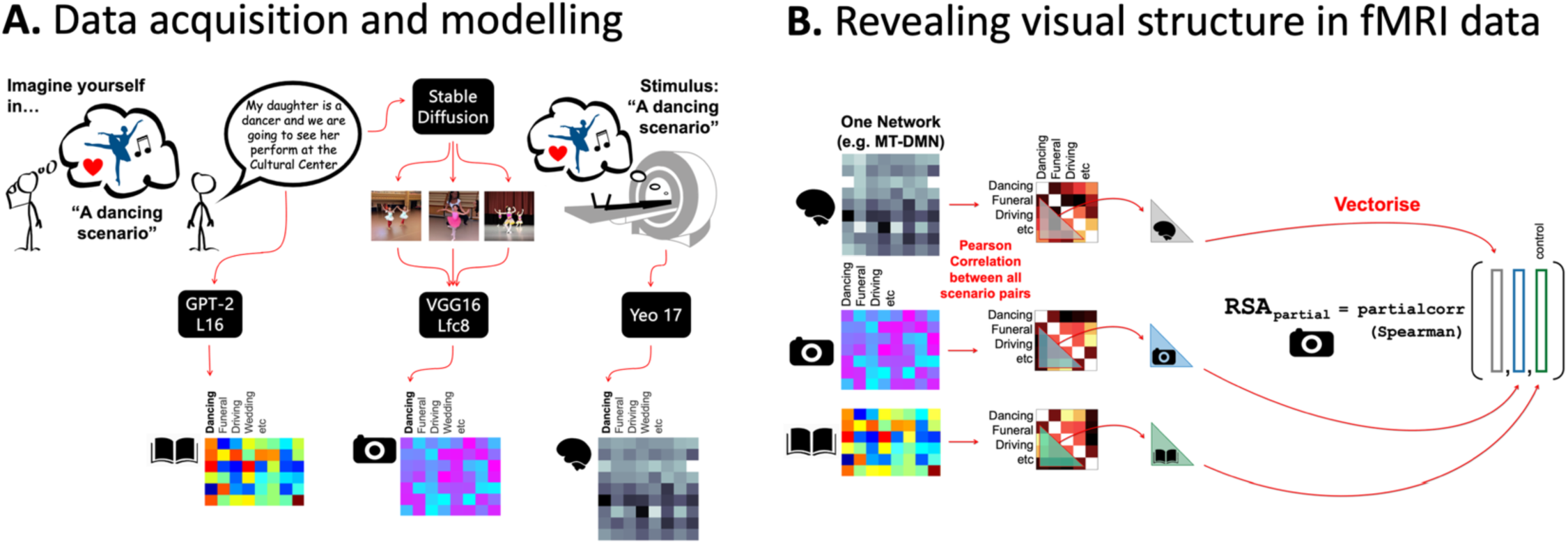
Identifying visual information structure in fMRI data scanned during autobiographical imagination. **A.** Before scanning, participants were presented with 20 generic scenario cues (e.g., dancing, exercising, wedding) and asked to vividly imagine themselves personally experiencing each event. They then verbally described each imagined scenario. During fMRI, participants re-imagined each scenario, presented in randomized order, based on generic text prompts. One fMRI volume was extracted per scenario per participant following standard preprocessing. To model visual scene structure, Stable Diffusion 2.1 (Rombach et al., 2022) was used to synthesize five images per scenario, guided by each participant’s description. The images were then passed through the object recognition network VGG-16 (Simonyan & Zisserman, 2015), and 1,000-dimensional activation vectors from the final pre-softmax layer (FC8) were extracted and averaged across the five images to yield a single embedding per scenario per participant (the Visual Model indicated by a Camera Icon). To control for non-visual semantic content, each verbal description was also modeled using GPT-2 L16 embeddings (Radford et al., 2019, Book Icon). **B.** To test for visual representational structure in the brain, fMRI patterns were extracted from the medial temporal default mode network (MT-DMN), as defined by the Yeo et al. (2011) parcellation. A partial correlation-based representational similarity analysis (RSA) was used to isolate variance explained uniquely by the Visual Model, controlling for GPT-2 embeddings. To test for specificity to MT-DMN, the analysis was repeated across Core- and Frontotemporal-DMN (FT-DMN) subdivisions, as well as the 14 remaining Yeo networks. In a complementary analysis, the models were switched in the partial RSA to examine variance explained uniquely by GPT-2 while controlling for the Visual Model.

Despite variability in viewpoint and object selection across different images for the same description, scene content and spatial layout were generally preserved in a contextually appropriate manner (e.g., people sitting on chairs rather than chairs on people). Many items and textures such as grass, tables, or bushes in a barbecue scene were inferred by the model even when not explicitly mentioned, reflecting the model’s learned priors about typical scene structure. Besides distortions, implausible content was rare.

To construct a participant-specific Visual Model suitable for comparison with fMRI data, each of the five images generated per scenario was processed through VGG-16, a convolutional neural network trained for object recognition (Simonyan & Zisserman, 2015). From each image, we extracted the 1,000-dimensional vector from the final pre-softmax layer (FC8), which encodes the model’s inferred likelihood of the presence of 1000 semantic categories, including animals, objects, buildings, landscape features, and other entities, based on the visual features of the input image (see **Methods**). The five FC8 vectors corresponding to the different images generated for each scenario were pointwise averaged to obtain a single, image embedding per scenario, per participant. Averaging increased robustness to occasional spurious image generations, and this approach proved more effective for downstream neural modeling than using a single image, as evaluated in subsequent analyses.

VGG-16 was selected for three main reasons. First, it has been widely used to model the neural correlates of object perception and imagery in humans (Nonaka et al., 2021), with related networks trained on ImageNet outperforming alternative approaches in predicting scene-viewing fMRI data (Groen et al., 2018). Second, the recognition features learned by VGG-16, including their divergence from human perceptual strategies, have been extensively characterized (Geirhos et al., 2019; Jacob et al., 2021; Bowers et al., 2023), offering interpretability benefits. Third, its output representations (FC8) were well-suited for averaging across multiple generated images, accommodating variability in spatial layout, viewpoint, and object composition that was not constant in the generated images, nor explicitly specified in participants’ scenario descriptions.

Because imagination is not exclusively visual in nature (e.g. Sulfaro et al. 2024), we deployed the Large Language Model GPT-2 (Radford et al. 2019) to model non-visual semantics associated with participants’ mental image descriptions. GPT-2 was selected because it is an established approach that has yielded strong results in predicting fMRI activation associated with natural language comprehension (Sun et al. 2020, Schrimpf et al. 2021, Caucheteux et al. 2022). Each mental image description was modeled with a single vector, derived by processing the entire description through GPT-2-medium, and pointwise averaging Layer 16 vector embeddings for all constituent tokens, following evidence that layers at 2/3 depth are appropriate (Caucheteux et al. 2022).

However, because the images synthesized by Stable Diffusion were driven by a textual semantic input representation derived from a separate text encoder (OpenClip or Open Contrastive Language–Image Pre-training, Cherti et al. 2023, see **Methods**), and OpenCLIP might potentially have explained Visual models findings more effectively than GPT-2, we ran supporting analyses to evaluate and ultimately rule out this possibility. Likewise, supporting analyses were run to determine whether re-representing the synthesized images with VGG-16 was a necessary step, and whether equivalent outcomes were possible using latent image representations extracted from within the Stable Diffusion model (See **Methods**).

### MT-DMN selectively reflected visual model representational structure

To test whether fMRI activity within the MT-DMN reflected visual representational structure, we applied a partial correlation-based representational similarity analysis (RSA; Kriegeskorte et al., 2008) to each participant’s data (Fig. 1B). This analysis evaluated whether the cluster structure of fMRI patterns across the 20 imagined scenarios aligned with that of the Visual Model, beyond GPT-2.

For example, different scenarios involving different people (e.g., family gathering or meeting a stranger) may share common visual features associated with human figures in social settings, that would be captured by the Visual Model. By contrast, GPT-2 may reflect differences in abstract semantic content between scenarios associated with familiarity and emotion valences associated with family and strangers. The partial correlation RSA was therefore designed to isolate variance in neural similarity patterns uniquely attributable to visual scene features, controlling for high-level non-visual semantic similarities.

To identify whether any visual effect detected was specific to MT-DMN, the same partial correlation RSA was repeated in Frontotemporal (FT) and Core-DMN which are also strongly linked to self-generated imagination/thought, and more specifically, to abstract semantic, verbal and social cognition (FT-DMN), and self-referential cognition (Core-DMN) (Andrews-Hanna and Grilli 2021, Andrews-Hanna et al. 2014, Andrews-Hanna et al. 2010, Alam et al. 2025). For completeness, fourteen other brain networks were also analyzed, that were predefined by the Yeo et al. (2011) parcellation scheme. To estimate the statistical significance associated with the set of partial RSA coefficients in each brain network, one sample signed ranks tests were run to compare correlation coefficients to zero (n=50 participants). Signed-rank p-values were adjusted across the seventeen networks according to False Discovery Rate (Benjamini and Hochberg, 1995).

To reveal non-visual semantic information structure, the procedure and group-level statistical tests were repeated, but with partial correlation RSA computed against GPT-2, while controlling for the Visual model. As context, the raw RSA coefficients between the Visual model and GPT-2 were Mean±SD: 0.21±0.09.

In support of the main hypothesis, MT-DMN selectively exhibited a strong sensitivity to the Visual Model comparative to the other sixteen networks (Fig. 2). MT-DMN’s visual selectivity was evaluated with one-tailed signed ranks tests, comparing the fifty MT-DMN Visual Model partial RSA coefficients to the corresponding coefficients from each of the other sixteen networks. Statistically significant outcomes were observed for all sixteen networks. The least significant outcome was for the Temporoparietal network (TempPar, Z=2.53, p=0.006), and Core-DMN (Z=2.55, p=0.005).

**Figure 2.**
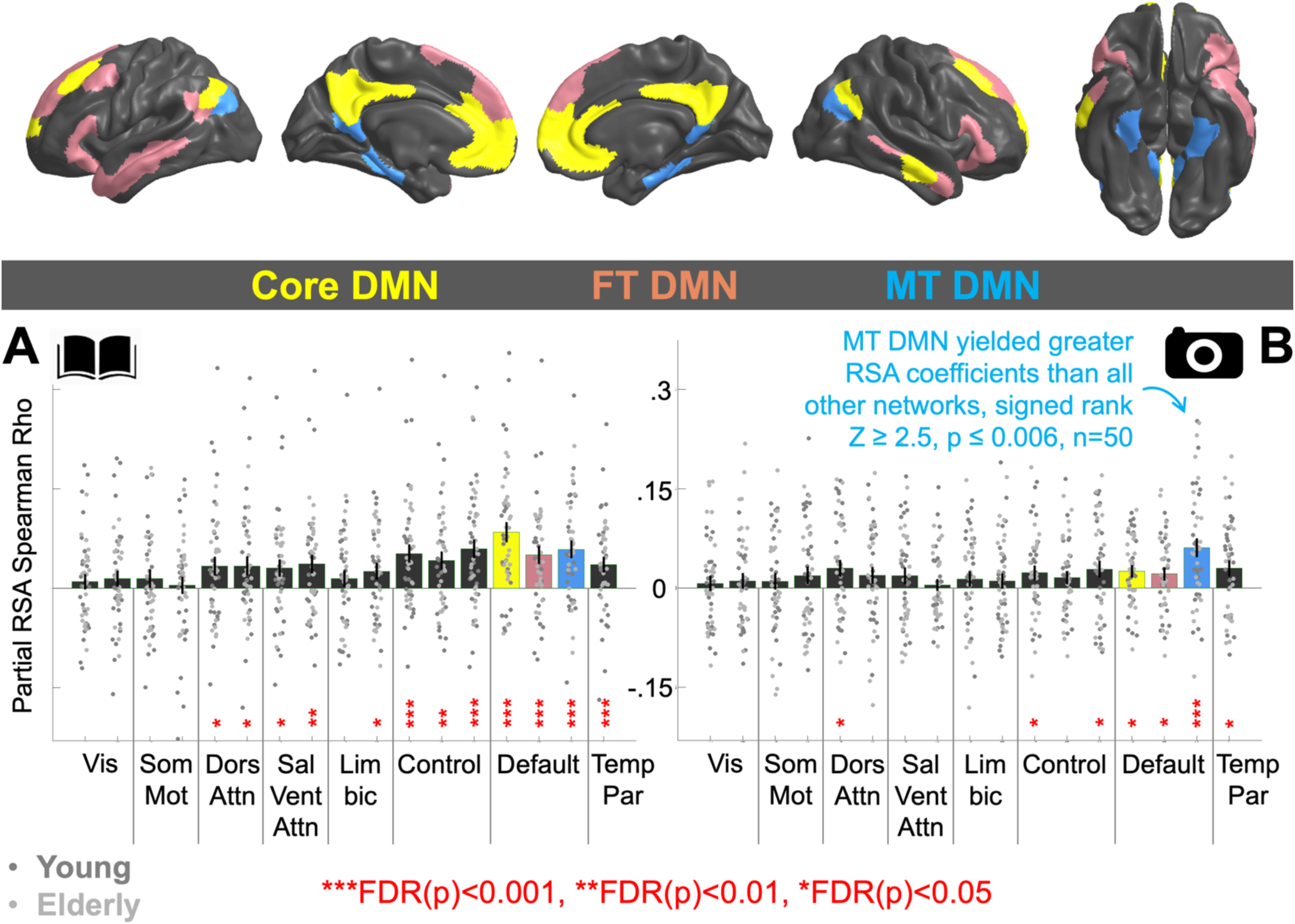
MT-DMN selectively reflects visual model representational structure. Partial correlation-based representational similarity analysis (Fig. 1A) was used to evaluate the unique contributions of visual and language models to fMRI pattern similarity across 17 brain networks from the Yeo et al. (2011) parcellation. Partial RSA coefficients are shown for each participant (N = 50). **A.** corresponds to GPT-2 (book icon). **B.** corresponds to the Visual Model derived from Stable Diffusion and VGG-16 (camera icon). In both plots, mean coefficients are plotted with error bars indicating ±1 SEM. P-values reflect signed-rank tests against zero, adjusted according to false discovery rate (Benjamini and Hochberg, 1995). The three default mode network (DMN) subsystems, medial temporal (MT-DMN), core (Core-DMN), and frontotemporal (FT-DMN), are highlighted with color-codes due to their hypothesized role in self-generated thought. The remaining 14 non-DMN networks are shown in grey to avoid distraction. Consistent with the overarching hypothesis, MT-DMN showed significantly greater sensitivity to the Visual Model than all other networks (Z ≥ 2.5, p ≤ 0.006, n=50, one tail). In contrast, GPT-2 not only explained similarity structure in MT-DMN but also in Core-DMN, where RSA coefficients were greater (Z = 2.6, p = 0.004, two tail, n=50), as well as eleven other networks (incl. FT-DMN). This suggests a broader encoding of non-visual semantic structure. **Supplementary** Fig. 1 reports RSA results without partial correlation. **Supplementary** Fig. 2 shows consistent findings when analyses were run separately in young and older adults. **Supplementary** Fig. 3 shows weaker outcomes when only one Stable Diffusion image, rather than five, was used. **Supplementary** Fig. 4 replicates findings using OpenCLIP in place of GPT-2. **Supplementary** Fig. 5 shows that latent representations from Stable Diffusion account for less variance in MT-DMN than the Visual Model used here. Brain visualizations were generated using BrainSpace v1.10 (Vos de Wael et al., 2020).

MT-DMN was also sensitive to GPT-2, but unlike the Visual model, this effect was not-specific. Core-DMN yielded significantly greater GPT-2 partial RSA coefficients than MT-DMN (signed-rank Z=2.6, p=0.004, two-tailed, n=50), and there were no significant differences between MT-DMN and FT-DMN, or the three Control networks.

Post hoc analyses demonstrated that: (1) The pattern of results in Fig. 2 broadly generalized to both young and elderly adults (**Supplementary** Fig. 2). (2) Basing the Visual model on five rather than one synthetic image was advantageous. (3) The Visual Model advantage in MT-DMN was replicated even more saliently when replacing GPT-2 with uOpenCLIP (Stable Diffusion’s textual input encoding model) (**Supplementary** Fig. 4). (4) Latent representations extracted from within Stable Diffusion were less accurate models of MT-DMN than the Visual model (**Supplementary** Fig. 5). (5) Fig. 2 could be replicated when images were resynthesized using a different implementation of Stable Diffusion (sdxl-turbo, Podell et al. 2023, **Supplementary** Fig. 6).

### MT-DMN selectively reflected participant-specific visual model structure

To assess whether MT-DMN encodes participant-specific visual structure stemming from an individual’s autobiographical experience, rather than more general semantics shared across participants, we performed a set of individual-differences analyses. These analyses aimed to discriminate participant identity by cross-correlating each participant’s fMRI data with their own and others’ participant-specific models.

Specifically, we compared the accuracy with which participants could be discriminated using: (1) A combined model incorporating the Visual Model with GPT-2. (2) GPT-2 alone. If the combined model discriminated participants’ fMRI patterns more effectively than GPT-2 alone, this would suggest that visual representations unique to the individual contributed to the neural signal, over and above what could be explained by participant-specific non-visual semantic structure.

To create the combined model, we first computed representational similarity matrices for each model (Visual and GPT-2), then vectorized and z-scored the lower triangle (Fig. 1B) of each matrix (mean = 0, SD = 1). The normalized vectors were then pointwise summed to form a composite matrix triangle. These combined similarity representations were then used in RSA to evaluate model fit to each participant’s fMRI data.

To evaluate whether fMRI data contained participant-specific visual information, we conducted a two-alternative forced-choice identity decoding analysis, following the method used in Anderson et al. (2020). This analysis tested how reliably pairs of participants could be distinguished based on how well their fMRI data matched their own versus another participant’s computational model.

For each pair of participants (P1 and P2), we computed the summed RSA fit for the congruent pairing: [RSA(fMRIP1,ModelP1)+RSA(fMRIP2,ModelP2)] and compared it to the incongruent pairing: [RSA(fMRIP1,ModelP2)+ RSA(fMRIP2,ModelP1)]. A decoding was counted as correct when the congruent sum was greater than the incongruent sum.

Under chance performance, correct discriminations would occur in 50% of comparisons. To assess significance, we used permutation testing, shuffling participant identities of the models but not fMRI data, and recalculating decoding accuracy across 10,000 iterations. A p-value was assigned based on the number of permutations in which the shuffled model outperformed the real model. P-values were corrected for multiple comparisons using False Discovery Rate (Benjamini and Hochberg, 1995).

As illustrated in Fig. 3A, participant identity was reliably discriminated by the Visual model in MT-DMN [72%, FDR(p) = 0.001] and, to a lesser extent, in FT-DMN [65%, FDR(p)=0.04]. In contrast, GPT-2 discriminated participant identity in 11 of the 17 networks, including MT-DMN [69%, FDR(p)=0.003), FT-DMN [66%, FDR(p) = 0.007], and Core-DMN [71%, FDR(p)=0.001]. These findings were replicated in both young and older adults (**Supplementary** Fig. 5).

**Figure 3.**
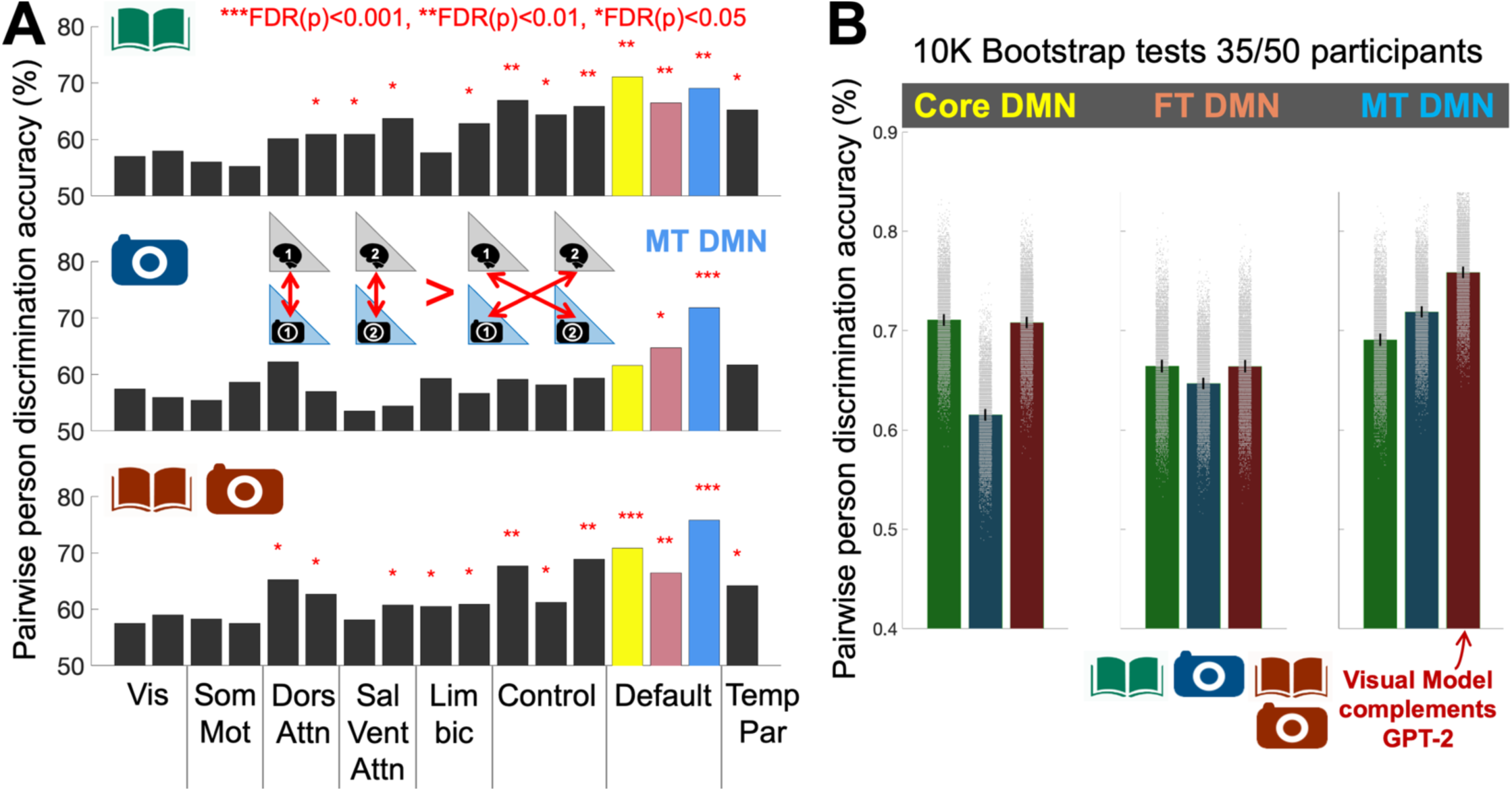
MT-DMN reflects participant-specific visual model structure. **A.** Individual-differences associated with the Visual model (Camera Icon) were prominent in MT-DMN and present in FT-DMN, whereas GPT-2 (Book Icon) captured individual-differences in 11/17 networks, including MT-, FT-, and Core-DMN. **B.** In support of the hypothesis that MT-DMN encodes participant-specific visual imagery, individual-differences in MT-DMN were more accurately discriminated when the Visual model and GPT-2 were combined (by averaging similarity vectors), than when GPT-2 was used in isolation (Red, Right Plot). The same effect was not observed in Core- and FT-DMN.

In MT-DMN, discrimination accuracy was highest when the Visual model and GPT-2 were combined [76%, FDR(p) = 0.0003], suggesting that the Visual model made an independent contribution to person-specific decoding. To evaluate this statistically, we ran a bootstrap analysis comparing the combined model to each isolated model. Specifically, the identity decoding analysis in Fig. 3A was repeated 10,000 times using random subsets of 35 out of 50 participants, and we counted how often the combined model yielded greater decoding accuracy than either isolated model.

Supporting the hypothesis that MT-DMN encodes participant-specific visual content, the combined model outperformed GPT-2 alone in 99.25% of bootstrapped samples (p = 0.0075, Fig. 3B). There was weaker evidence that GPT-2 made an independent contribution: the combined model outperformed the Visual model in only 91% of subsamples.

For comparison, this analysis was repeated in FT-DMN and Core-DMN. The only other significant difference emerged in Core-DMN, where participant-specific decoding was driven entirely by GPT-2: GPT-2 alone outperformed the Visual model in 99.9% of subsamples and showed comparable performance to the combined model (GPT-2 > combined in 56% of subsamples).

### MT-DMN did not reflect visual model structure during sentence reading without active imagination

Finally, to verify that the visual structure detected in MT-DMN (Figure 2) was a product of imagination, as opposed to a more general property of semantic representation, a control analysis was run on a separate fMRI dataset scanned as 14 separate people read 240 sentences without any imagination task (Anderson et al. 2017b). The 240 sentences all described scenes and/or objects, such as: “The girl broke the glass at the shop” (**Supplementary Table 1**). The sentences were modeled using Stable Diffusion/GPT-2 using precisely the same approach as the autobiographical scenario descriptions (Fig. 1A) and the fMRI data was analyzed using same partial correlation RSA approach (**Fig. 1B** and the same group-level statistical tests (Fig. 2) on the same seventeen cortical networks (Yeo et al. 2011).

Consistent with the hypothesis that the visual information structure previously detected in MT-DMN in Fig. 2**/3** stemmed from active imagination, MT-DMN did not reflect Visual model structure to a significant degree during sentence reading, without an imagination task (Fig. 4). The Visual model did however make a statistically significant contribution to explaining fMRI data structure in Core-DMN. However, the magnitude of this effect was very small comparative to GPT-2, which captured fMRI structure in all seventeen networks, with far greater correlation coefficients in all three DMN subsystems.

**Figure 4.**
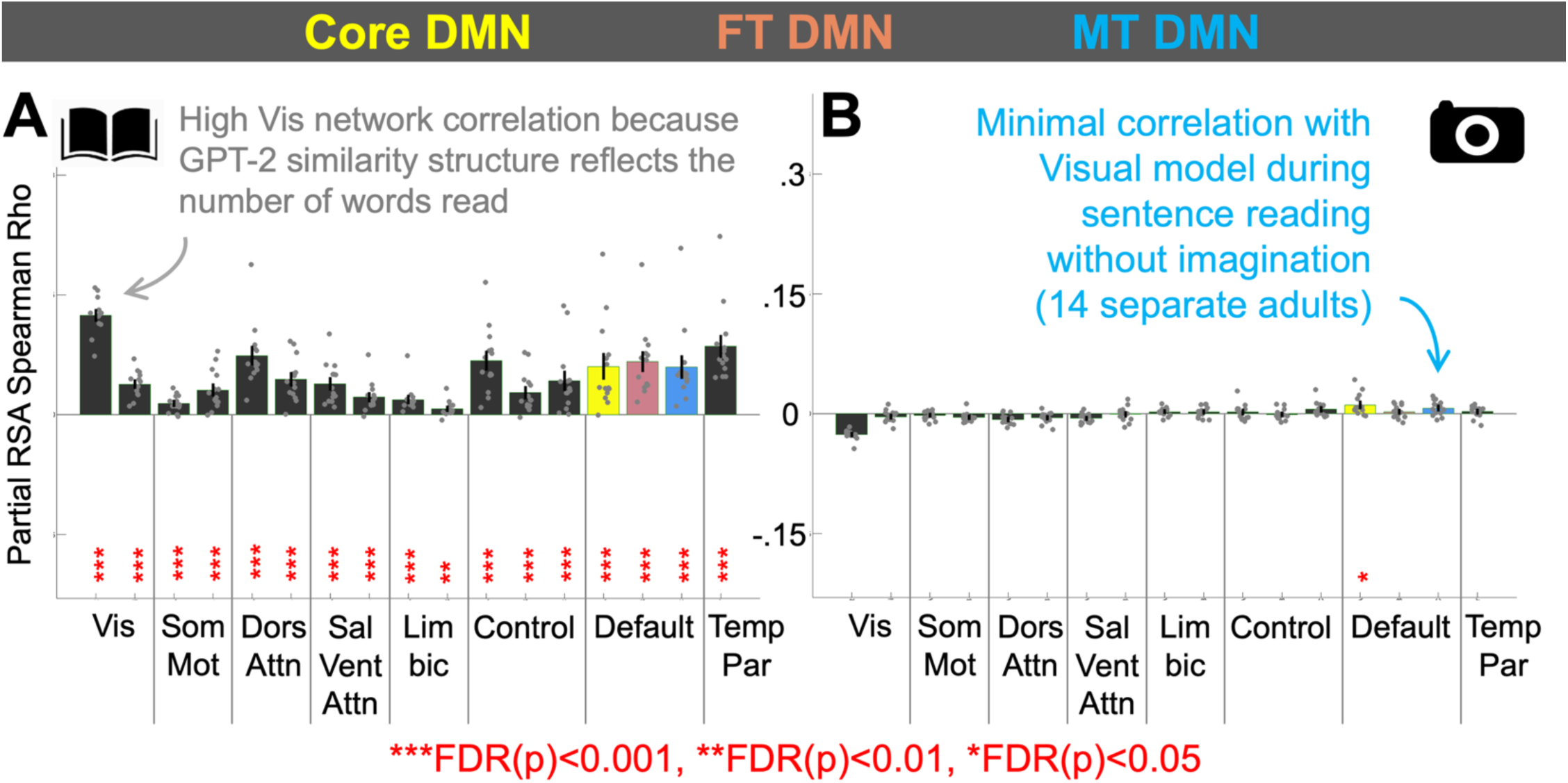
MT-DMN did not reflect visual structure during sentence reading without active imagination. To estimate whether the visual structure detected in MT-DMN in Fig. 2 was a product of active imagination, the analyses in Fig. 1 were repeated on a separate fMRI dataset scanned as fourteen different people read 240 simple sentences about events and objects, without any active imagination task (Anderson et al. 2017). In this case, MT-DMN did not reflect visual information structure, however a small statistically significant effect was observed in Core-DMN (Yellow). Differently, GPT-2 captured information structure across all seventeen brain networks. This included an especially strong effect in “Vis” networks which are most associated with active vision, as opposed to language semantics. This same effect has been observed in previous analysis of the same data (Anderson et al. 2021) and stems from correlations between the representational similarity structure associated with sentence/word-lengths and language model embeddings. Bars indicate mean RSA coefficients across participants. Error bars are SEM. P-values correspond to signed-ranks tests computed across all fifty people, adjusted according to False Discovery Rate (Benjamini and Hochberg, 1995).

## Discussion

This study provides evidence that fMRI activation patterns within the medial temporal default mode network selectively reflect participant-specific visual reconstructions of autobiographical experiences, generated in response to generic textual prompts. Critically, such representations were not observed in other large-scale networks, nor during a sentence reading task where mental imagery was not required. These findings support the role of the MT-DMN in reconstructing visual scenes from memory (Andrews-Hanna & Grilli, 2021; Andrews-Hanna et al., 2010, 2014), thereby helping to delineate its function relative to other brain networks. The study also provides proof-of-principle that freely available image generation (Rombach et al., 2022; Podell et al., 2023) and image recognition systems (Simonyan & Zisserman, 2015) can be combined to build individualized brain models of imagined visual scenes derived from participants’ verbal self-reports.

In doing so, the study bridges three major areas of research: (1) Self-generated imagery and thought, which has often relied on coarse experience sampling to model subjective mental content (e.g. Delamillieure et al., 2010; Gorgolewski et al., 2014; Smallwood & Schooler, 2015; Sormaz et al., 2018); (2) Episodic recall and prospection, which have identified brain representations of distinct events (Chadwick et al., 2010; Baldassano et al., 2017; Chen et al., 2017), and encoded information about time, place, people, and objects (Bonnici et al., 2012; Nielson et al., 2015; Thornton et al., 2017; Silson et al., 2019; Sreekumar et al., 2018; Benoit et al., 2019; Gagnepain et al., 2020; Anderson et al., 2020; Gilmore et al., 2021); (3) Visual imagery, where neural representations of isolated objects have been modeled using image-based computational approaches (Reddy et al., 2010; Cichy et al., 2012; Ragni et al., 2020; Anderson et al., 2015; Soto et al., 2020; Horikawa & Kamitani, 2017; Breedlove et al., 2020). However, unlike prior studies, the current approach introduces a method to discriminate participant-specific visual representations of autobiographical scenes. This distinction enabled us to isolate idiosyncratic autobiographical visual imagery within MT-DMN, a capability not previously demonstrated to our knowledge.

The current findings suggest that the MT-DMN generates visual correlates of semantic categories in scene contexts. To some degree these visual correlates reflect the image statistics exploited by VGG-16 to classify scene contents generated by Stable Diffusion, from participant text. While further work is needed to determine which aspects of this visual pipeline were most critical, we tentatively attribute results to two key factors. First, Stable Diffusion may enrich scene reconstructions by inferring visual contextual details, even when these are not explicitly mentioned in text prompts. For example, a “barbecue scene” might be visually rendered with food on a table on a lawn, despite these features, and their arrangement being omitted from the prompt. Such enrichment, though occasionally inaccurate, may generally be advantageous in reintroducing visual details that participants considered too obvious to report, and on average may better approximate mental images. Second, VGG-16 may leverage contextual and textural information, beyond just object shape or silhouette, when producing image embeddings, and this may capture related contextual representations in scene selective cortices (Bar, 2003, 2004; Aminoff et al., 2013) that overlap MT-DMN. This assertion follows evidence that classification in image recognition models can rely heavily on background context, such that a cat-shaped object with an elephant skin texture is recognized as an elephant and not a cat (Geirhos et al., 2019), and a hatchet is only be recognized in a woodland, but not when it is displayed amongst vegetables in a grocery store (Jacob et al., 2021). It is also consistent with studies showing that image recognition models (Groen et al. 2018) more accurately predict neural responses than manually annotated lists of object categories within images (c.f. Stansbury et al., 2013; Çukur et al., 2016).

The ability to model neural representations associated with relatively freeform imagination may have applications for understanding individual differences across cognitive domains and clinical populations. Mental imagery capacity varies widely, from individuals with aphantasia - who report no voluntary visual imagery - to those with hyperphantasia, who describe imagery as vivid as perception (Galton, 1880; Zeman, 2024; Dawes et al., 2020). These differences are not only of theoretical interest but may also inform how people retrieve, construct, or distort internal experiences such as autobiographical memories or future simulations. Mental imagery also plays a central role in numerous psychological disorders. For example, involuntary, emotionally charged imagery is a hallmark of flashbacks in post-traumatic stress disorder, vivid craving in addiction, and mood-congruent imagery in bipolar disorder (Pearson et al., 2015; Holmes & Mathews, 2010). In schizophrenia and Parkinson’s disease, patients often report heightened or altered sensory imagery (Aleman et al., 2005; Fénelon et al., 2000). Conversely, individuals with depression may experience a diminished ability to imagine positive future scenarios (Ji et al., 2016), while those with suicidal ideation often report vivid flash-forwards to suicidal actions (Holmes et al., 2007). By capturing participant-specific visual representations, the current framework may offer a path toward quantifying and comparing the neural substrates of imagery across such diverse conditions.

While the current visual model appears useful for modeling representations within the MT-DMN, it is important to emphasize that this does not imply that the brain constructs such representations in the same manner. Our approach applies two transformations, going from synthesized images and then to recognition embeddings, akin to applying perceptual analysis to imagination. However, this contrasts with evidence suggesting that mental imagery may operate in the reverse direction, whereby stored high-level visual representations are projected backward along the visual hierarchy to reconstruct low-level sensory details in early visual cortex (Pearson et al., 2019; Dijkstra et al., 2020; Breedlove et al., 2020). In principle, deriving embeddings directly from the latent layers of Stable Diffusion could better align with the generative direction of imagination. However, in practice, our attempts to use latent embeddings from within Stable Diffusion were less effective than those produced by VGG-16 (**Supplementary** Figures 4,5). Nonetheless, improving generative model alignment with brain-based reconstruction processes remains a promising direction for future work.

Finally, a second limitation involves spatial structure. Prior studies indicate that imagined scenes retain spatial layout and viewpoint information (Hassabis et al., 2007a, 2007b; Hassabis & Maguire, 2007; Andrews-Hanna & Grilli, 2021; Alam et al., 2025). However, such details were often underspecified in our participants’ verbal descriptions and therefore were unlikely to be fully recovered during image generation or even preserved in 2D image embeddings. Future work could address this by collecting detailed spatial descriptions or sketch-based inputs and reconstructing three-dimensional models from the resulting images to better capture scene layout and viewpoint.

In conclusion, this study demonstrates that the medial temporal default mode network makes a unique contribution to autobiographical visual imagery, as identified through modeling fMRI data with participant-specific visual reconstructions of verbal self-reports. This approach offers a framework for studying freeform imagination in the human brain and may provide a useful tool for characterizing individual differences in mental imagery, particularly in contexts where imagery is exaggerated, diminished, or atypical.

## Methods

### Autobiographical Mental Imagery Participants

50 participants’ data were reanalyzed. 50% of the data was collected from healthy young adults (Mean±SD age=24±3, 16F, 9M, Wang et al. 2024). The other 25 were healthy elderly adults (Mean±SD age was 73±7 years, 16F, 9M, Anderson et al. 2020). One elderly participant (#26) was excluded from the current study because of discordant BOLD/structural fMRI co-registration when the data were preprocessed with fMRIPrep for the current study (SPM had been used in Anderson et al. 2020).

### Autobiographical Mental Imagery Scenario Stimuli

Twenty scenarios were preselected by Anderson et al. 2020 to be diverse events that almost every participant would have personally experienced. Scenarios were always presented to participants in the following form: “A X Scenario”, or “An X Scenario” where X is a placeholder for: resting, reading, writing, bathing, cooking, housework, exercising, internet, telephoning, driving, shopping, movie, museum, restaurant, barbecue, party, dancing, wedding, funeral, festival.

### Autobiographical Mental Imagery Experimental Procedure

At the beginning of the experiment, an experimenter went through the twenty scenario prompts asking participants to vividly imagine themselves experiencing each scenario, and to actively simulate their sensory experiences, actions and feelings. Once participants had formed a rich mental image, they provided a brief verbal description of their mental image which was transcribed by the experimenter. After all the scenarios had been imagined and described, the experimenter went through the scenarios again, first reminding the participant of their scenario description, and then recording the participant’s ratings of the likelihood that their mental image reflected a real personal experience (rather than being fiction) and the vividness of the mental image (see **Supplementary Table 1**).

### Autobiographical Mental Imagery MRI Data Collection Parameters

Imaging data were collected at the University of Rochester Center for Brain Imaging using a 3T Siemens Prisma scanner (Erlangen, Germany) equipped with a 32-channel receive-only head coil. The fMRI scan began with a MPRAGE scan (TR/TE=1400/2344 ms, TI=702ms, Flip Angle=8°, FOV=256mm, matrix=256×256mm, 192 sagittal slices, slice thickness =1mm, voxel size 1×1×1mm3). fMRI data were collected using a gradient echo-planar imaging (EPI) sequence (TR/TE=2500ms/30ms, Flip Angle=85°, FOV=256mm, 90 axial slices, slice thickness=2mm, voxel size 2×2×2mm3, number of volumes = 639).

### Autobiographical Mental Imagery fMRI Experiment

Prior to scanning, participants were reminded of their verbal descriptions of the 20 scenarios (see above) and were requested to vividly reimagine the same mental images in the scanner, when prompted. Outside the scanner they then underwent a single dry run of the fMRI experiment below (a single viewing of all 20 stimuli) to familiarize them with the set up.

In the experiment, stimulus prompts (e.g., “A dancing scenario”) were presented one by one in random order, on a screen in black Arial font (size 50) on a grey background. Prompts remained on screen for 7.5 seconds (3TRs), during which time participants vividly imaged themselves in the scenario. After the prompt was removed there was a 7.5 second delay prior to the next prompt (e.g., “A barbecue scenario”) during which time a fixation cross was displayed. Participants had been instructed to attempt to clear their minds when the prompts were removed.

Participants then underwent a single uninterrupted fMRI session in which the 20 scenario stimuli were presented five times over (five runs). The five repeats later enabled us to compute representations of average mental images (per scenario) and counteract fMRI noise. Stimulus order was randomized within each run. Runs were separated by a 15-second interval, in which a second-by-second countdown was displayed (e.g., “Starting run 2 in 13 seconds”), which was followed by 7.5 seconds of fixation cross preceding the first stimulus of the run. All participants reported that they had been able to imagine the scenarios on prompt.

### Depicting participants’ textual descriptions of autobiographical mental images with Stable Diffusion

Stable Diffusion (Rombach et al., 2022; Podell et al., 2023) is a text-to-image generative model that synthesizes realistic images from random noise, guided by a natural language prompt. In the current study, these prompts were participants’ verbal descriptions of autobiographical mental images. The generation process is carried out through reverse diffusion, which iteratively denoises a latent image representation based on the semantic embedding of the text prompt.

This process proceeds in three main stages. First, the text prompt is encoded using OpenCLIP (Cherti et al., 2023), a vision-language model trained on image-caption pairs, producing a sequence of high-dimensional embedding vectors that capture the visual semantics of the described scene.

Second, reverse diffusion is applied in a compressed latent space (64 × 64 × 4) to generate a denoised image embedding, seeded with an embedding of noise. This is guided by the OpenCLIP embedding and reverses a pre-trained forward diffusion process that had learned to iteratively degrade images by adding noise. Finally, a trained autoencoder decodes the denoised latent representation into a 512 × 512 RGB image.

For this study, we used the MLX implementation of Stable Diffusion 2.1 on macOS (15.4.1, 24E263) with default parameters: 50 diffusion steps and a classifier-free guidance (CFG) scale of 7.5, which adjusts the fidelity of the image to the text prompt. For each of the twenty imagined scenarios described by each participant (N = 50), five distinct images were generated from different initial noise vectors.

### Deriving a Visual Model by processing Stable Diffusion Images with VGG-16

VGG-16 (Simonyan & Zisserman, 2015) is a convolutional neural network (CNN) trained on over one million images from the ImageNet database (Deng et al., 2009) to classify images into 1,000 categories, including buildings, environmental features, objects, animals, plants, and food. The network consists of 16 layers with learnable weights: 13 convolutional layers followed by three fully connected layers (Fc6, Fc7, Fc8). The Fc6 and Fc7 layers each contain 4,096 units, while Fc8 contains 1,000 units corresponding to the classification categories. These outputs are typically passed through a softmax function to produce class probabilities.

For the current analysis, each 512 × 512 RGB image generated by Stable Diffusion was resized to 224 × 224 and passed through VGG-16. We extracted the 1,000-dimensional vector from the final pre-softmax layer (Fc8) to represent the image, following prior fMRI and neural decoding studies (Horikawa & Kamitani, 2017; Nonaka et al., 2021; Anderson et al., 2017; Groen et al., 2018), as well as work in multimodal computational semantics (Kiela & Bottou, 2014).

For each participant and each autobiographical scenario, five images were synthesized using Stable Diffusion. The corresponding Fc8 vectors were averaged pointwise to produce a single 1,000-dimensional visual representation per scenario, per participant.

### Deriving a semantic control model of participants textual mental image descriptions with GPT-2

GPT-2 (Generative Pretrained Transformer 2, Radford et al. 2019 is a Transformer Decoder Deep-Learning Model trained entirely upon text to predict the identity of the next word, given a history of up to 1024 preceding words. GPT-2 is widely believed to induce representations that serve as a proxy for semantics, in internal layers, that help it to predict impending words. Given evidence that layers at ∼2/3 depth are a good choice for modeling natural language comprehension fMRI data (Caucheteux et al. 2022). Each mental image description was modeled with a single vector of 1024 features, derived by processing the sentence/description through GPT-2-medium and pointwise averaging Layer 16 (of 24) activation for all constituent tokens.

### Autobiographical Mental Imagery MRI Data Preprocessing

MRI data were preprocessed using fMRIPrep (Esteban et al. 2019) to counteract head motion and spatially normalize images to a common neuroanatomical space (MNI152NLin2009cAsym). The boilerplate template detailing the procedure is in **Supplementary Materials**. To counteract nuisance signals and potential confounds in the fMRI data a comprehensive selection of nuisance regressors generated by fMRIPrep were regressed out from each voxels time series. These were 24 head motion parameters (including translation, rotation and their derivatives), white-matter and cerebrospinal fluid timeseries and cosine00, cosine01, cosine02, cosine03. Confound removal was implemented through computing a single multiple regression across the entire fMRI timeline: First, each voxel and each nuisance regressor’s time-series was separately z-scored (by subtracting the mean and dividing by the standard deviation). Second a separate multiple regression was fit for each voxel, mapping nuisance regressors to predict voxel activation. The residuals from the regression (computed separately for each voxel) were taken forward to further analysis (below).

For analyses, we reduced each participant’s fMRI data to obtain a single volume for each imagined scenario. To do this we first computed a single volume for each scenario replicate, by computing the voxel-wise mean of 4 fMRI volumes (5-15sec) post stimulus onset. To reduce the five replicates of each scenario to a single scenario volume, we again took the voxel-wise mean. This left 20 scenario volumes per individual. The 5 (2TR) second offset was to accommodate hemodynamic response delay, so if a participant were to instantly bring their mental image to mind, the peak response would be around 4-5 seconds (in practice, mental images are unlikely to have been formed instantly). The four-volume period spans the time until the next stimulus is presented on screen and was set to maximize our chances of capturing the mental image. NB The current averaging approach was selected to accommodate individual differences in the latency and duration of hemodynamic responses associated with internally generated mental images observed by Anderson et al. 2020, which may not be well suited to modeling with a canonical hemodynamic response function time-locked to stimulus presentation.

### Sentence Comprehension Participants

Participants were 14 healthy, native speakers of English (5 males, 9 females; mean age = 32.5, range 21-55) with no history of neurological or psychiatric disorders. All were right-handed according to the Edinburgh Handedness Inventory (Oldfield 1971).

### Sentence Comprehension Stimuli

The stimuli, listed in full in **Supplementary Table 2**, consisted of 240 written sentences containing 3-9 words and 2-5 (Mean±SD = 3.33±.76) content words, formed from different combinations of 141 nouns, 62 verbs, and 39 adjectives (242 words). Sentences were in active voice and consisted of a noun phrase followed by a verb phrase in past tense, with no relative clauses. Most sentences (200/240) contained an action verb and involved interactions between humans, animals and objects, or described situations involving different entities, events, locations, and affective connotations. The remaining 40 sentences contained only a linking verb (“was”). All sentences were pre-selected as experimental materials for the Knowledge Representation in Neural Systems (KRNS) project (Glasgow et al. 2016, www.iarpa.gov/index.php/research-programs/krns), sponsored by the Intelligence Advanced Research Projects Activity (IARPA).

### Sentence Comprehension Experimental Procedure

Participants took part in multiple scanning visits, where they were instructed to read sentences and think about their overall meaning (as detailed in Anderson et al. 2017). In each visit, the entire list of sentences was presented 1.5 times in 12 scanning runs, with each run containing 30 trials (one sentence per trial) and lasting approximately 6 minutes. The presentation order of each set of 240 sentences was randomly shuffled. Sentences were presented word-by-word using a rapid serial visual presentation paradigm. Nouns, verbs, adjectives, and prepositions were presented for 400 ms each, followed by a 200ms inter-stimulus interval (ISI). Articles (“the”) were presented for 150 ms followed by a 50-ms ISI. Mean sentence duration was 2.8 s (range [1.4s 4.2s]). Words subtended an average horizontal visual angle of approximately 2.5°.

### Sentence Comprehension MRI parameters and preprocessing

MRI data were acquired with a whole-body 3T GE 750 scanner at the Center for Imaging Research of the Medical College of Wisconsin using a GE 32-channel head coil. Functional T2*-weighted echoplanar images (EPI) were collected with TR = 2000 ms, TE = 24 ms, flip angle = 77°, 41 axial slices, FOV = 192 mm, in-plane matrix = 64 x 64, slice thickness = 3 mm, resulting in 3 x 3 x 3 mm voxels. T1-weighted anatomical images were obtained using a 3D spoiled gradient-echo sequence with voxel dimensions of 1 x 1 x 1 mm.

### Sentence Comprehension MRI Data Preprocessing

fMRI data are publicly available (Anderson et al. 2021) and were preprocessed using AFNI (Cox, 1996) in Anderson et al. 2017. EPI volumes were corrected for slice acquisition time and head motion. Functional volumes were aligned to the T1-weighted anatomical volume, transformed into a standardized space (Talairach and Tournoux 1988), and smoothed with a 6mm FWHM Gaussian kernel. The data were analyzed using a general linear model with a duration-modulated HRF, and the model included one regressor for each sentence. fMRI activity was modeled as a gamma function convolved with a square wave with the same duration as the presentation of the sentence, as implemented in AFNI’s 3dDeconvolve with the option dmBLOCK. Duration was coded separately for each individual sentence. Finally, a single sentence-level fMRI representation was created for each unique sentence by taking the voxel-wise mean of all replicates of the sentence.

## Data and Code

Data and code to replicate analyses are available at DOI: 10.17605/OSF.IO/TM92D.

## References

Addis DR, Wong AT, Schacter, DL 2007. Remembering the past and imagining the future: common and distinct neural substrates during event construction and elaboration. Neuropsychologia 45, 1363–1377.

Alam TR, Krieger-Redwood K, Varga D, Gao Z, Horner AJ, Hartley T, de Schotten MT, Sliwinska M, Pitcher D, Margulies DS, Smallwood J. 2025. A double dissociation between semantic and spatial cognition in visual to default network pathways. Elife.13:RP94902.

Aleman A, Böcker KB, Hijman R, de Haan EH, Kahn RS. 2005. Cognitive basis of hallucinations in schizophrenia: Role of top–down information processing. Schizophrenia Research, 74(2–3), 233–241.

Aminoff EM, Kveraga K, Bar M. The role of the parahippocampal cortex in cognition. Trends in cognitive sciences. 2013 Aug 1;17(8):379–90.

Anderson AJ, Bruni E, Lopopolo A, Poesio M, Baroni M. 2015. Reading visually embodied meaning from the brain: Visually grounded computational models decode visual-object mental imagery induced by written text. NeuroImage. 120:309–22.

Anderson AJ, Binder JR, Fernandino L, Humphries CJ, Conant LL, Aguilar M, Wang X, Doko D, Raizada RDS. 2017. Predicting neural activity patterns associated with sentences using a neurobiologically motivated model of semantic representation. Cereb Cortex 27:4379–4395.

Anderson AJ, Kiela D, Clark S, Poesio M. 2017. Visually grounded and textual semantic models differentially decode brain activity associated with concrete and abstract nouns. Transactions of the Association for Computational Linguistics. 1;5:17–30.

Anderson AJ, McDermott K, Rooks B, Heffner KL, Dodell-Feder D, Lin FV. 2020. Decoding individual identity from brain activity elicited in imagining common experiences. Nature Communications. 20;11(1):1–4.

Anderson AJ, Kiela D, Binder JR, Fernandino L, Humphries CJ, Conant LL, Raizada RDS, Grimm S, Lalor EC. 2021. Deep artificial neural networks reveal a distributed cortical network encoding propositional sentence-level meaning. Journal of Neuroscience. 41(18):4100–19.

Andrews-Hanna JR, Reidler JS, Sepulcre J, Poulin R, Buckner RL. 2010. Functional-anatomic fractionation of the brain’s default network. Neuron. 65(4):550–62.

Andrews-Hanna JR, Smallwood J, Spreng RN. 2014. The default network and self-generated thought: Component processes, dynamic control, and clinical relevance. Annals of the New York Academy of Sciences. 1316(1):29–52.

Andrews-Hanna JR, Grilli MD. 2021. Mapping the imaginative mind: Charting new paths forward. Current Directions in Psychological Science. 30(1):82–9.

Axelrod V, Rees G, Bar M. 2017. The default network and the combination of cognitive processes that mediate self-generated thought. Nature Human Behaviour. 1(12):896–910.

Baldassano C, Chen J, Zadbood A, Pillow JW, Hasson U, Norman KA. 2017. Discovering event structure in continuous narrative perception and memory. Neuron. 95(3):709–21.

Bar M, Aminoff E. 2003. Cortical analysis of visual context. Neuron. 38(2):347–58.

Bar M. Visual objects in context. 2004. Nature Reviews Neuroscience. 5(8):617–29.

Benjamini Y, Hochberg Y. 1995. Controlling the false discovery rate: a practical and powerful approach to multiple testing. Journal of the Royal statistical society: series B (Methodological). 57(1):289–300.

Benoit RG, Paulus PC, Schacter DL. 2019. Forming attitudes via neural activity supporting affective episodic simulations. Nature communications. 10(1):1–1.

Bonnici HM, Chadwick MJ, Lutti A, Hassabis D, Weiskopf N, Maguire EA. 2012. Detecting representations of recent and remote autobiographical memories in vmPFC and hippocampus. Journal of Neuroscience. 32(47):16982–91.

Breedlove JL, St-Yves G, Olman CA, Naselaris T. 2020. Generative feedback explains distinct brain activity codes for seen and mental images. Current Biology. 30(12):2211–24.

Bowers JS, Malhotra G, Dujmović M, Montero ML, Tsvetkov C, Biscione V, Puebla G, Adolfi F, Hummel JE, Heaton RF, Evans BD. 2023. Deep problems with neural network models of human vision. Behavioral and Brain Sciences. 46:e385.

Buckner RL, DiNicola LM. 2019. The brain’s default network: updated anatomy, physiology and evolving insights. Nature Reviews Neuroscience. 20(10):593–608.

Cant JS, Xu Y. 2012. Object ensemble processing in human anterior-medial ventral visual cortex. J. Neurosci. 32:7685–700

Caucheteux C, Gramfort A, King J-R. 2022. Deep language algorithms predict semantic comprehension from brain activity. Scientific reports 12.1 16327.

Chadwick MJ, Hassabis D, Weiskopf N, Maguire EA. 2010. Decoding individual episodic memory traces in the human hippocampus. Current Biology. 20(6):544–7.

Cherti M, Beaumont R, Wightman R, Wortsman M, Ilharco G, Gordon C, Schuhmann C, Schmidt L, Jitsev J; Proceedings of the IEEE/CVF Conference on Computer Vision and Pattern Recognition (CVPR), 2023, pp. 2818–2829.

Chen J, Leong YC, Honey CJ, Yong CH, Norman KA, Hasson U. 2017. Shared memories reveal shared structure in neural activity across individuals. Nature Neuroscience. 20(1):115–25.

Cichy RM, Heinzle J, Haynes JD. 2012. Imagery and perception share cortical representations of content and location. Cerebral cortex. 22(2):372–80.

Çukur T, Huth AG, Nishimoto S, Gallant JL. 2016. Functional subdomains within scene-selective cortex: parahippocampal place area, retrosplenial complex, and occipital place area. The Journal of Neuroscience 36: 10257–10273.

Dawes AJ, Keogh R, Andrillon T, Pearson J. 2020. A cognitive profile of multi-sensory imagery, memory and dreaming in aphantasia. Scientific Reports, 10, 10022.

Delamillieure P, Doucet G, Mazoyer B, Turbelin MR, Delcroix N, Mellet E, Zago L, Crivello F, Petit L, Tzourio-Mazoyer N, Joliot M. 2010. The resting state questionnaire: An introspective questionnaire for evaluation of inner experience during the conscious resting state. Brain research bulletin. 81(6):565–73.

Deng J, Dong W, Socher R, Li L-J, Li K, Fei-Fei L. 2009. Imagenet: A large-scale hierarchical image database. In: 2009 IEEE conference on computer vision and pattern recognition. p. 248–55.

Dijkstra N, Ambrogioni L, Vidaurre D, van Gerven M. 2020. Neural dynamics of perceptual inference and its reversal during imagery. e2020 Jul 20;9:e53588.

Dijkstra N, Bosch SE, van Gerven MA. 2019. Shared neural mechanisms of visual perception and imagery. Trends in cognitive sciences. 23(5):423–34.

Eickenberg M, Gramfort A, Varoquaux G, Thirion B. 2017. Seeing it all: Convolutional network layers map the function of the human visual system. NeuroImage. 152:184–94.

Epstein RA, Baker CI. 2019. Scene perception in the human brain. Annual review of vision science. 5(1):373–97.

Esteban O, Markiewicz CJ, Blair RW, Moodie CA, Isik AI, Erramuzpe A, Kent JD, Goncalves M, DuPre E, Snyder M, Oya H, Ghosh SS, Wright J, Durnez J, Poldrack RA, Gorgolewski KJ. 2019. fMRIPrep: a robust preprocessing pipeline for functional MRI. Nat Methods. 16(1):111–116. doi: 10.1038/s41592-018-0235-4.

Fénelon G, Mahieux F, Huon R, Ziegler M. 2000. Hallucinations in Parkinson’s disease: Prevalence, phenomenology and risk factors. Brain, 123(4), 733–745.

Fernandino L, Binder JR. 2024. How does the “default mode” network contribute to semantic cognition? Brain and Language. 252, 105405

Galton, F. Statistics of mental imagery. 1880. Mind 5, 301–318.

Gagnepain P, Vallée T, Heiden S, Decorde M, Gauvain JL, Laurent A, Klein-Peschanski C, Viader F, Peschanski D, Eustache F. 2020. Collective memory shapes the organization of individual memories in the medial prefrontal cortex. Nature Human Behaviour. 4(2):189–200.

Geirhos R, Rubisch P, Michaelis C, Bethge M, Wichmann FA, Brendel W. 2019. ImageNet-trained CNNs are biased towards texture; increasing shape bias improves accuracy and robustness. In International Conference on Learning Representations (ICLR), https://arxiv.org/abs/1811.12231

Gilmore AW, Quach A, Kalinowski SE, Gotts SJ, Schacter DL, Martin A. 2021. Dynamic content reactivation supports naturalistic autobiographical recall in humans. Journal of Neuroscience. 2021 41(1):153–66.

Glasgow K, Roos M, Haufler A, Chevillet M, Wolmetz, M, 2016. Evaluating semantic models with word-sentence relatedness. arXiv:1603.07253.

Gong Z, Zhou M, Dai Y, Wen Y, Liu Y, Zhen Z. 2023. A large-scale fMRI dataset for the visual processing of naturalistic scenes. Scientific Data. 10(1):559.

Gorgolewski KJ, Lurie D, Urchs S, Kipping JA, Craddock RC, Milham MP, Margulies DS, Smallwood J. 2014. A correspondence between individual differences in the brain’s intrinsic functional architecture and the content and form of self-generated thoughts. PloS one. 9(5):e97176.

Groen IIA, Greene MR, Baldassano C, Li FF, Beck DM, Baker CI. 2018. Distinct contributions of functional and deep neural network features to representational similarity of scenes in human brain and behavior. eLife 7:e32962

Hassabis, D., Kumaran, D., Vann, S. D. & Maguire, E. A. Patients with hippocampal amnesia cannot imagine new experiences. Proc. Natl Acad. Sci. USA 104, 1726–1731 (2007).

Hassabis, D., Kumaran, D. & Maguire, E. A. Using imagination to understand the neural basis of episodic memory. J. Neurosci. 27, 14365–14374 (2007).

Hassabis D, Maguire EA. Deconstructing episodic memory with construction. 2007. Trends in Cognitive Sciences. 11:299–306

Ji JL, Holmes EA, MacLeod C, Murphy, FC. 2016. Mental imagery in psychiatry: Conceptual and clinical implications. CNS Spectrums, 21(4), 335–340.

Holmes EA, Mathews A. 2010. Mental imagery in emotion and emotional disorders. Clinical Psychology Review, 30(3), 349–362.

Holmes EA, Crane C, Fennell MJV, Williams JMG. 2007. Imagery about suicide in depression - “Flash-forwards”? Journal of Behavior Therapy and Experimental Psychiatry, 38(4), 423–434.

Horikawa T, Kamitani Y. 2017 Generic decoding of seen and imagined objects using hierarchical visual features. Nature communications. 8.1: 15037.

Horikawa T. 2025. Mind captioning: Evolving descriptive text of mental content from human brain activity. Science Advances. 11, 45 DOI: 10.1126/sciadv.adw146

Jacob G, Pramod RT, Katti H, Arun SP. 2021. Qualitative similarities and differences in visual object representations between brains and deep networks. Nature communications.12(1):1872.

Johnson MR, Johnson MK. 2014. Decoding individual natural scene representations during perception and imagery. Frontiers in human neuroscience. 8:59.

Khaligh-Razavi SM, Kriegeskorte N. 2014. Deep supervised, but not unsupervised, models may explain IT cortical representation. PLoS computational biology. 10(11):e1003915.

Kiela D, Bottou L. 2014. Learning image embeddings using convolutional neural networks for improved multi-modal semantics. In Proceedings of EMNLP, pages 36–45, Doha, Qatar.

Kriegeskorte N, Mur M, Bandettini PA. 2008. Representational similarity analysis-connecting the branches of systems neuroscience. Frontiers in systems neuroscience. 24;2:249.

Leeds DD, Seibert DA, Pyles JA, Tarr MJ. 2013. Comparing visual representations across human fMRI and computational vision J. Vis., 13 (13), pp. 1–27 (25)

Lowe DG. 2004. Distinctive image features from scale-invariant keypoints. International journal of computer vision. 60:91–110.

Mars RB, Neubert FX, Noonan MP, Sallet J, Toni I, Rushworth MF. 2012. On the relationship between the “default mode network” and the “social brain”. Frontiers in human neuroscience. 6:189.

Matsuyama T, Nishimoto S, Takagi Y. et al. 2025.LaVCa: LLM-assisted Visual Cortex Captioning. arXiv preprint arXiv:2502.13606.

Menon V. 2023. 20 years of the default mode network: A review and synthesis. Neuron. 111(16):2469–87.

Nielson DM, Smith TA, Sreekumar V, Dennis S, Sederberg PB. 2015. Human hippocampus represents space and time during retrieval of real-world memories. Proceedings of the National Academy of Sciences. 112(35):11078–83.

Nishida S, Nishimoto S. 2018. Decoding naturalistic experiences from human brain activity via distributed representations of words. Neuroimage. 180:232–42.

Nishimoto S, Vu AT, Naselaris T, Benjamini Y, Yu B, Gallant JL. 2011. Reconstructing visual experiences from brain activity evoked by natural movies. Current Biology. 21(19):1641–6.

Nonaka S, Majima K, Aoki SC, Kamitani Y. 2021. Brain hierarchy score: Which deep neural networks are hierarchically brain-like?. IScience. 24(9).

O’Craven KM, Kanwisher N. 2000. Mental imagery of faces and places activates corresponding stimulus-specific brain regions. Journal of cognitive neuroscience. 12(6):1013–23.

Oldfield RC. 1971. The assessment and analysis of handedness: The Edinburgh Inventory. Neuropsychologia. (9):97–113.

Pearson J, Naselaris T, Holmes EA, Kosslyn SM. 2015. Mental imagery: Functional mechanisms and clinical applications. Trends in Cognitive Sciences. 19(10), 590–602.

Pearson J. 2019. The human imagination: the cognitive neuroscience of visual mental imagery. Nature reviews Neuroscience. 20(10):624–34.

Podell D, English Z, Lacey K, Blattmann A, Dockhorn T, Müller J, Penna J, Rombach R. 2023.Sdxl: Improving latent diffusion models for high-resolution image synthesis. arXiv preprint arXiv:2307.01952.

Radford A, Wu J, Child R, Luan D, Amodei D, Sutskever I. 2019. Language models are unsupervised multitask learners. OpenAI blog, 1(8):9.

Ragni F, Tucciarelli R, Andersson P, Lingnau A. 2020. Decoding stimulus identity in occipital, parietal and inferotemporal cortices during visual mental imagery. Cortex. 1;127:371–87.

Raichle ME, MacLeod AM, Snyder AZ, Powers WJ, Gusnard DA, Shulman GL. 2001. A default mode of brain function. Proceedings of the National Academy of Sciences. 98(2):676–82.

Reddy L, Tsuchiya N, Serre T. 2010. Reading the mind’s eye: decoding category information during mental imagery. Neuroimage. 50(2):818–25.

Robin J, Buchsbaum BR, Moscovitch M. 2018. The primacy of spatial context in the neural representation of events. Journal of Neuroscience. 38(11):2755–65.

Robinson AK, Quek GL, Carlson TA. 2023. Visual representations: Insights from neural decoding. Annual Review of Vision Science. 9(1):313–35.

Rombach R, Blattmann A, Lorenz D, Esser P, Ommer B. 2022. High-resolution image synthesis with latent diffusion models. In Proceedings of the IEEE/CVF conference on computer vision and pattern recognition 2022 (pp. 10684–10695).

Schacter DL and Addis DR. 2007. The ghosts of past and future. Nature 445, 27–27.

Schacter DL, Addis DR, Hassabis D, Martin VC, Spreng RN, Szpunar KK. 2012. The future of memory: remembering, imagining, and the brain. Neuron. 76(4):677–94.

Schacter DL, Addis DR, Buckner RL. 2007. Remembering the past to imagine the future: the prospective brain. Nat. Rev. Neurosci. 8, 657–661.

Schaefer A, Kong R, Gordon EM, Laumann TO, Zuo XN, Holmes AJ, Eickhoff SB, Yeo BTT. 2018. Local-Global parcellation of the human cerebral cortex from intrinsic functional connectivity MRI. Cerebral Cortex, 29:3095–3114.

Schrimpf M, Blank IA, Tuckute G, Kauf C, Hosseini EA, Kanwisher N, Tenenbaum JB, Fedorenko E. The neural architecture of language: Integrative modeling converges on predictive processing. Proceedings of the National Academy of Sciences. 118(45):e2105646118.

Shao X, Krieger-Redwood K, Zhang M, Hoffman P, Lanzoni L, Leech R, Smallwood J, Jefferies E. 2024. Distinctive and complementary roles of default mode network subsystems in semantic cognition. Journal of Neuroscience. 44(20).

Silson EH, Steel A, Kidder A, Gilmore AW, Baker CI. 2019. Distinct subdivisions of human medial parietal cortex support recollection of people and places. Elife. 8.

Simony E, Honey CJ, Chen J, Lositsky O, Yeshurun Y, Wiesel A, Hasson U. 2016. Dynamic reconfiguration of the default mode network during narrative comprehension. Nature Communications. 7(1):12141.

Simonyan K, Zisserman A. Very deep convolutional networks for large-scale image recognition. 2014. arXiv preprint arXiv:1409.1556.

Smallwood, J, Schooler, JW. 2015. The science of mind wandering: empirically navigating the stream of consciousness. Annu. Rev. Psychol. 66, 3487–3518.

Smallwood J, Bernhardt BC, Leech R, Bzdok D, Jefferies E, Margulies DS. 2021. The default mode network in cognition: a topographical perspective. Nature reviews neuroscience. 22(8):503–13.

Sormaz M, Murphy C, Wang HT, Hymers M, Karapanagiotidis T, Poerio G, Margulies DS, Jefferies E, Smallwood J. 2018. Default mode network can support the level of detail in experience during active task states. Proceedings of the National Academy of Sciences. 115(37):9318–23.

Soto D, Sheikh UA, Mei N, Santana R. 2020. Decoding and encoding models reveal the role of mental simulation in the brain representation of meaning. Royal Society open science. 20;7(5):192043.

Spreng RN, Mar RA, Kim AS. 2009. The common neural basis of autobiographical memory, prospection, navigation, theory of mind, and the default mode: a quantitative meta-analysis. Journal of Cognitive Neuroscience. 21(3):489–510.

Sreekumar V, Nielson DM, Smith TA, Dennis SJ, Sederberg PB. 2018. The experience of vivid autobiographical reminiscence is supported by subjective content representations in the precuneus. Scientific reports. 8(1):1–9.

Stansbury DE, Naselaris T, Gallant JL. 2013. Natural scene statistics account for the representation of scene categories in human visual cortex. Neuron 79:1025–34

Sulfaro AA, Robinson AK, Carlson TA. Properties of imagined experience across visual, auditory, and other sensory modalities. Consciousness and Cognition. 2024 Jan 1;117:103598.

Sun J, Wang S, Zhang J, Zong C. 2020. Neural encoding and decoding with distributed sentence representations. IEEE Transactions on Neural Networks and Learning Systems. 32(2):589–603.

Szpunar KK, Watson JM, McDermott KB. Neural substrates of envisioning the future. 2007. Proc. Natl Acad. Sci. USA 9, 104.

Tang J, LeBel A, Jain S, Huth AG. 2023. Semantic reconstruction of continuous language from non-invasive brain recordings. Nature Neuroscience. 1:1–9.

Thornton MA, Mitchell JP. 2017. Consistent neural activity patterns represent personally familiar people. Journal of cognitive neuroscience. (9):1583–94.

Tulving, E. 1972. Episodic and semantic memory. In Organization of Memory, E. Tulving and W. Donaldson, eds. (Academic Press), pp. 381–403.

Ungerleider LG, Mishkin M. 1982. Two cortical visual systems. In: Ingle, D.J.,Goodale, M.A., Mansfield, R.J.W. (Eds.), Analysis of Visual Behavior. MIT Press, Cambridge, MA, pp. 549–586.

Ungerleider LG, Haxby JV. 1994. ‘What’ and ‘where’ in the human brain. Curr. Opin. Neurobiol. 4, 157–165.

Wang Y, Turnbull A, Xiang T, Xu Y, Zhou S, Masoud A, Azizi S, Lin FV, Adeli E. 2024. Decoding Visual Experience and Mapping Semantics through Whole-Brain Analysis Using fMRI Foundation Models. arXiv preprint arXiv:2411.07121.

Wheeler MA, Stuss DT, Tulving E. 1997. Toward a theory of episodic memory: the frontal lobes and autonoetic consciousness. Psychological bulletin. 121(3):331.

Yamins DL, Hong H, Cadieu CF, Solomon EA, Seibert D, DiCarlo JJ. Performance-optimized hierarchical models predict neural responses in higher visual cortex. Proceedings of the national academy of sciences. 2014 Jun 10;111(23):8619–24.

Yeo BT, Krienen FM, Sepulcre J, Sabuncu MR, Lashkari D, Hollinshead M, Roffman JL, Smoller JW, Zöllei L, Polimeni JR, Fischl B. 2011. The organization of the human cerebral cortex estimated by intrinsic functional connectivity. Journal of neurophysiology. 106(3):1125.

Yeshurun Y, Nguyen M, Hasson U. 2021. The default mode network: Where the idiosyncratic self meets the shared social world. Nature Reviews Neuroscience, 22(3), 181–192.

Zeman A. 2024. Aphantasia and hyperphantasia: exploring imagery vividness extremes. Trends in Cognitive Sciences. 28(5):467–80.

