## Supplementary Materials for "Medial Temporal Default Mode Network Selectively Encodes Autobiographical Visual Imagery"

### Contents:

**Supplementary Figure 1.** RSA coefficients corresponding to the Visual Model, GPT-2, and their combination 2

**Supplementary Figure 2.** MT-DMN selectively reflects visual content in both young and elderly adults 3

**Supplementary Figure 3.** Repeating **Fig. 2** using only one synthetic image rather than five yields a weaker outcome 4

**Supplementary Figure 4. Fig. 2** outcomes are replicated when OpenCLIP is used in place of GPT-2, against the Visual model (Stable Diffusion and VGG-16[Fc8]) 5

**Supplementary Figure 5.** Repeating **Fig. 2** with Stable Diffusion Latent Representations in place of the Visual model (Stable Diffusion and VGG-16[Fc8]) yields weaker MT-DMN sensitivity 6

**Supplementary Figure 6. Fig. 2** results are replicated with a different Stable Diffusion model (Stable Diffusion XL Turbo) 7

**Supplementary Figure 7.** MT-DMN reflects participant-specific visual content during imagination in young and elderly adults 8

**Supplementary Table 1.** Experimental sentences. 9

**fMRIPrep Boilerplate Template** 14

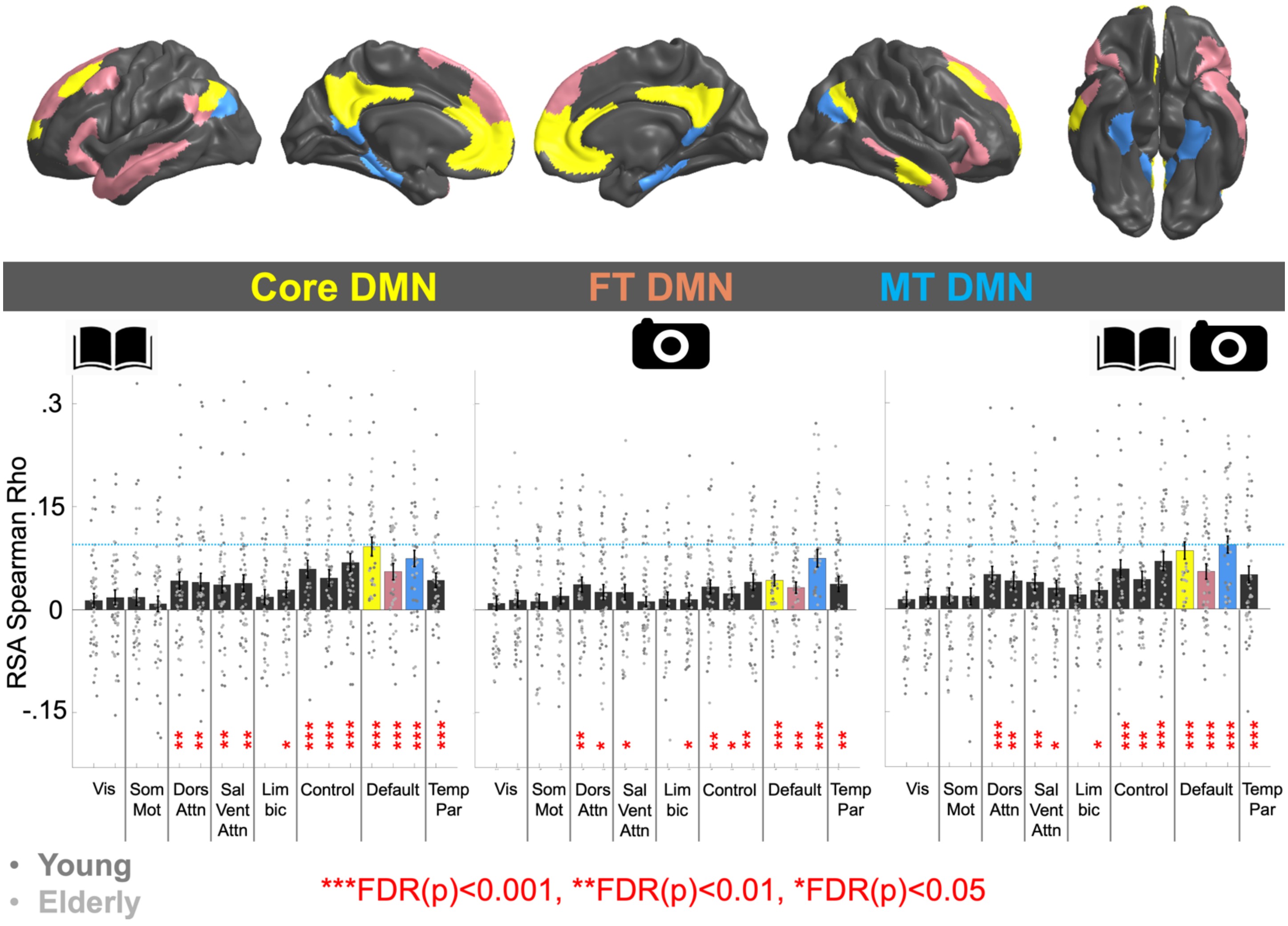


**Supplementary Figure 1.** RSA coefficients corresponding to the Visual Model, GPT-2, and their combination

Raw RSA coefficients computed using correlation against isolated models are displayed for fifty participants in each of the seventeen Yeo networks, to support the partial correlation-based results displayed in **Fig. 2**. GPT-2 (Book icon) is Left, the Visual model (Camera Icon) is Middle, and the two models combined (Book and Camera Icon) are Right. Model combination was achieved at the RSA stage by pointwise averaging together the two correlation matrix triangles (see **Fig. 1 Step 2**), after each triangle was normalized by z-scoring (mean zero, SD=1). Results for the three DMN subsystems are color coded. Bars indicate mean RSA coefficients across participants. Error bars are SEM. P-values correspond to signed-ranks tests computed across all fifty people, adjusted according to False Discovery Rate (FDR, Benjamini and Hochberg, 1995).

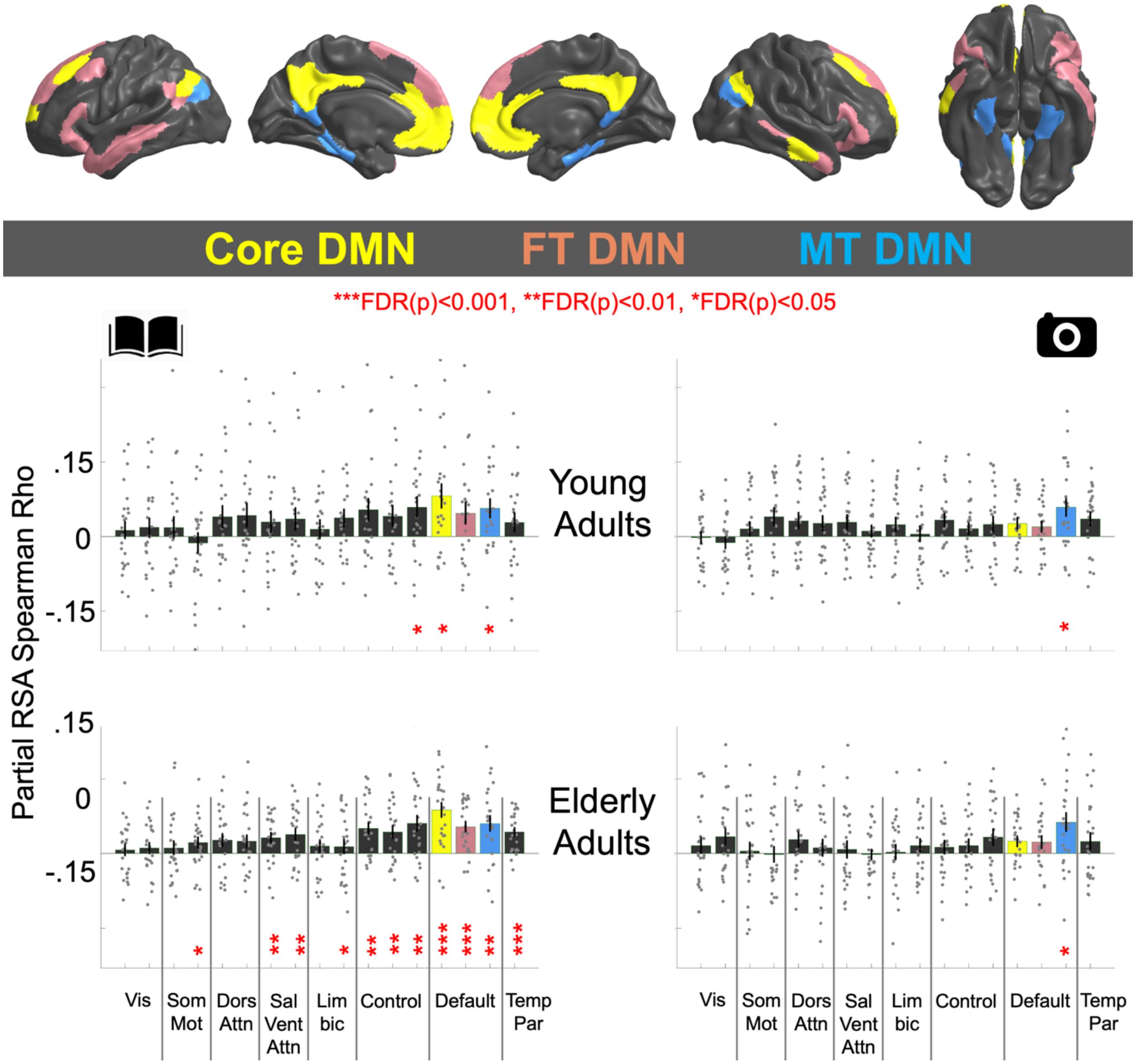


**Supplementary Figure 2.** MT-DMN selectively reflects visual content in both young and elderly adults

Partial correlation RSA coefficients (**Figure 1 Step 2**) corresponding to GPT-2 (Left, Book icon) and the Visual model (Right, Camera Icon) are displayed for 25 young adult participants and 25 elderly adult participants in each of the seventeen Yeo networks. The three DMN subsystems are color coded. MT-DMN selectively reflected the Visual model with statistical significance in both young and elderly adults. In young adults the set of 25 Visual model partial correlation coefficients in MT-DMN was significantly greater than fourteen of the sixteen networks, with the least significant differences observed for the Temporoparietal network (Z=1.08, p=0.14, one-tail) and Somatosensory-motor network B (Z=1.24, p=0.11, one-tail). In elderly adults, the set of 25 Visual model partial correlation coefficients in MT-DMN was significantly greater than fifteen of the sixteen other networks, with the least significant difference observed for the Peripheral Visual network (Z=1.5, p=0.07, one-tail). GPT-2 also made a strong contribution to explaining MT-DMN, but also contributed to other networks, especially Core-DMN in young and elderly adults. Bars indicate mean RSA partial coefficients across participants. Error bars are SEM. P-values correspond to signed-ranks tests computed across all fifty people, adjusted according to False Discovery Rate (Benjamini and Hochberg, 1995). GPT-2 also made a strong contribution to MT-DMN, but this was not-specific to MT-DMN, with Core-DMN yielding significantly greater partial correlation coefficients (signed-rank Z=2.6, p=0.004), and no significant differences for FT-DMN, and the three Control networks.

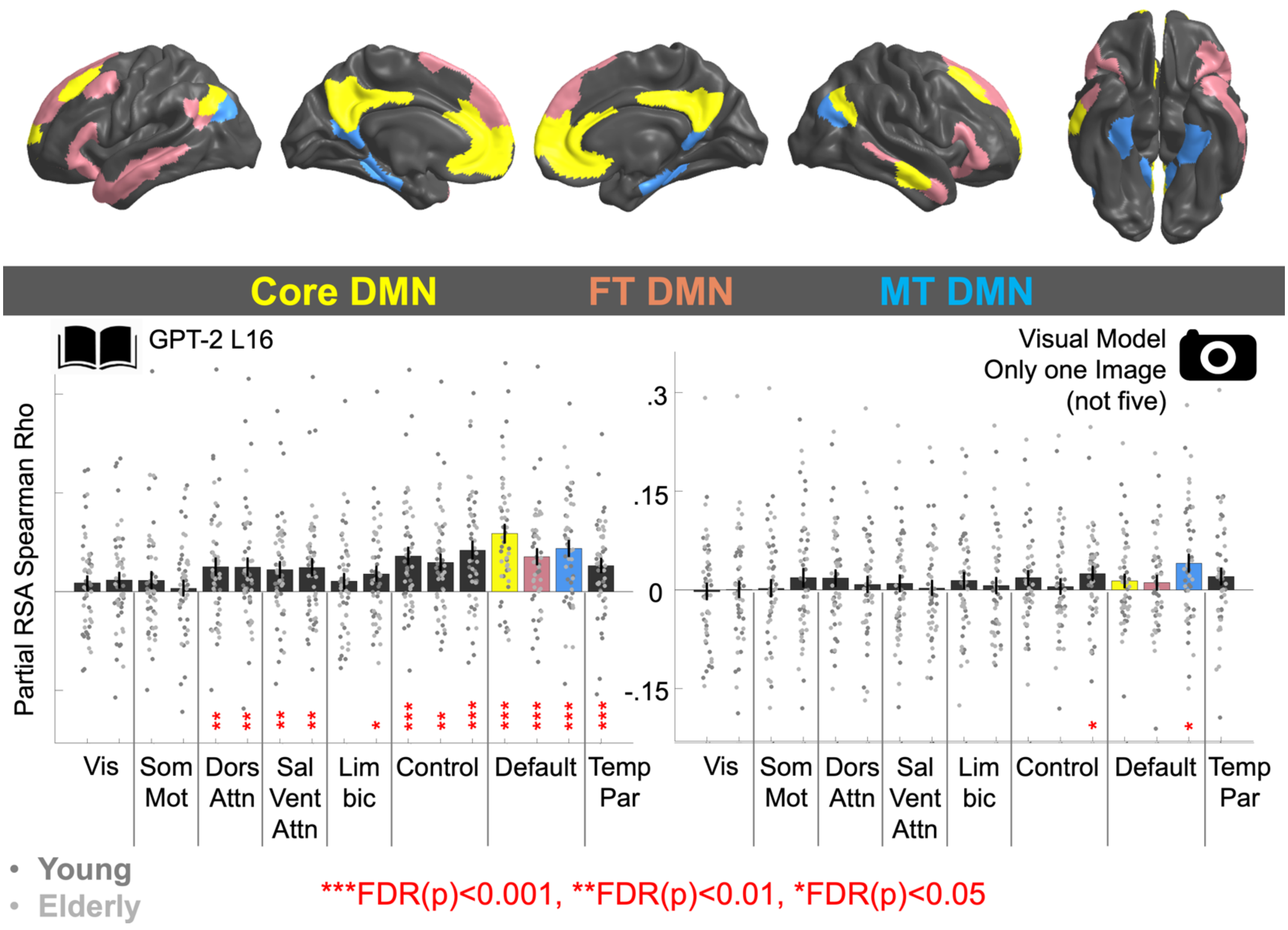


**Supplementary Figure 3.** Repeating **Fig. 2** using only one synthetic image rather than five yields a weaker outcome

To evaluate whether generating five synthetic images for each imagined autobiographical scenario description was valuable - as opposed to generating one single image (which might be less robust to spurious outcomes in image generation), the analyses of **Fig .2** were repeated based on only one synthetic image per scenario. Partial correlation RSA coefficients (**Figure 1 Step 2**) corresponding to GPT-2 (Left, Book icon) and the Visual model (Right, Camera Icon) are displayed, when the Visual model was constructed using only a single synthetic image rather than five (**Fig. 2**). Values are displayed for fifty participants in each of the seventeen Yeo networks. Visual model Partial RSA coefficients in MT-DMN were significantly lower when only one image was synthesized rather then five images (**Fig. 2**), z=2.1, p=0.035 (2-tail), n=50. Nonetheless, MT-DMN still exhibited statistically greater sensitivity to the Visual model than fourteen of the other networks, besides Control C (z=1.3, p=0.01) and TempPar z=1.5, p=0.06).

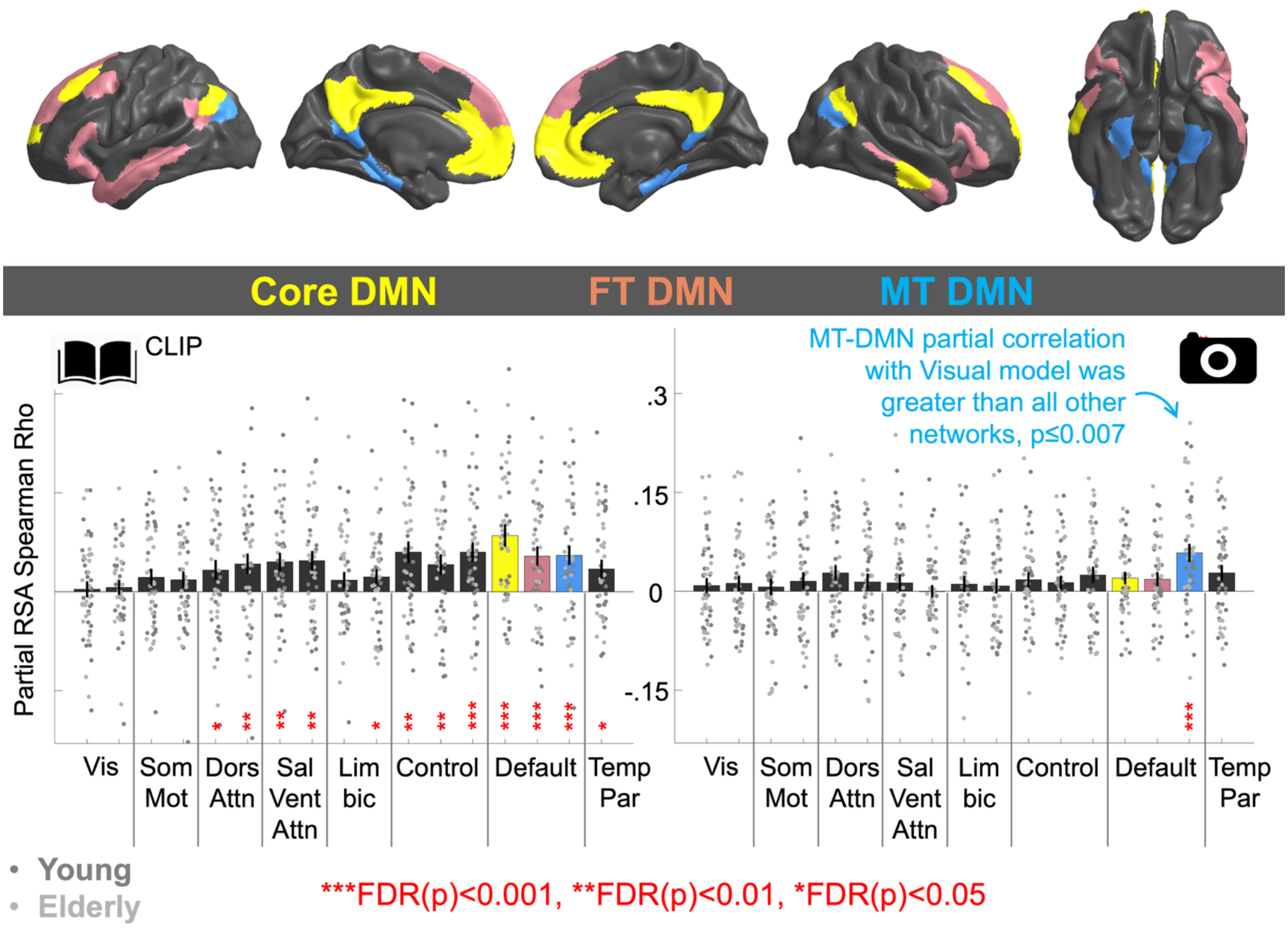


**Supplementary Figure 4. Fig. 2** outcomes are replicated when OpenCLIP is used in place of GPT-2, against the Visual model (Stable Diffusion and VGG-16[Fc8])

To evaluate whether the text-based OpenCLIP (Cherti et al. 2023) representation input to guide image generation in Stable Diffusion 2.1 captured activation patterns that had been attributed to the Visual model in **Fig. 2**, the analyses of **Fig .2** were repeated using OpenCLIP in place of GPT-2. Partial correlation RSA coefficients derived from OpenCLIP, when controlling for the Visual model are displayed in the Left plot (Book Icon). The Right plot (Camera Icon) displays partial correlation RSA coefficients derived from the Visual model, when controlling for OpenCLIP. Critically, MT-DMN alone was strongly captured by the Visual model. MT-DMN's selective sensitivity was statistically evaluated with one-tailed signed ranks tests, comparing the fifty MT-DMN Visual model partial correlation coefficients to corresponding coefficients from each of the other sixteen networks. Statistically significant outcomes were observed for all sixteen networks, with the least significant outcomes in Temporoparietal network (TempPar, Z=2.46, p=0.007), and Core-DMN (2.51, p=0.006). Bars indicate mean RSA partial correlation coefficients across participants. Error bars are SEM. P-values correspond to signed-ranks tests against zero, computed across all fifty people, adjusted according to False Discovery Rate (Benjamini and Hochberg, 1995).

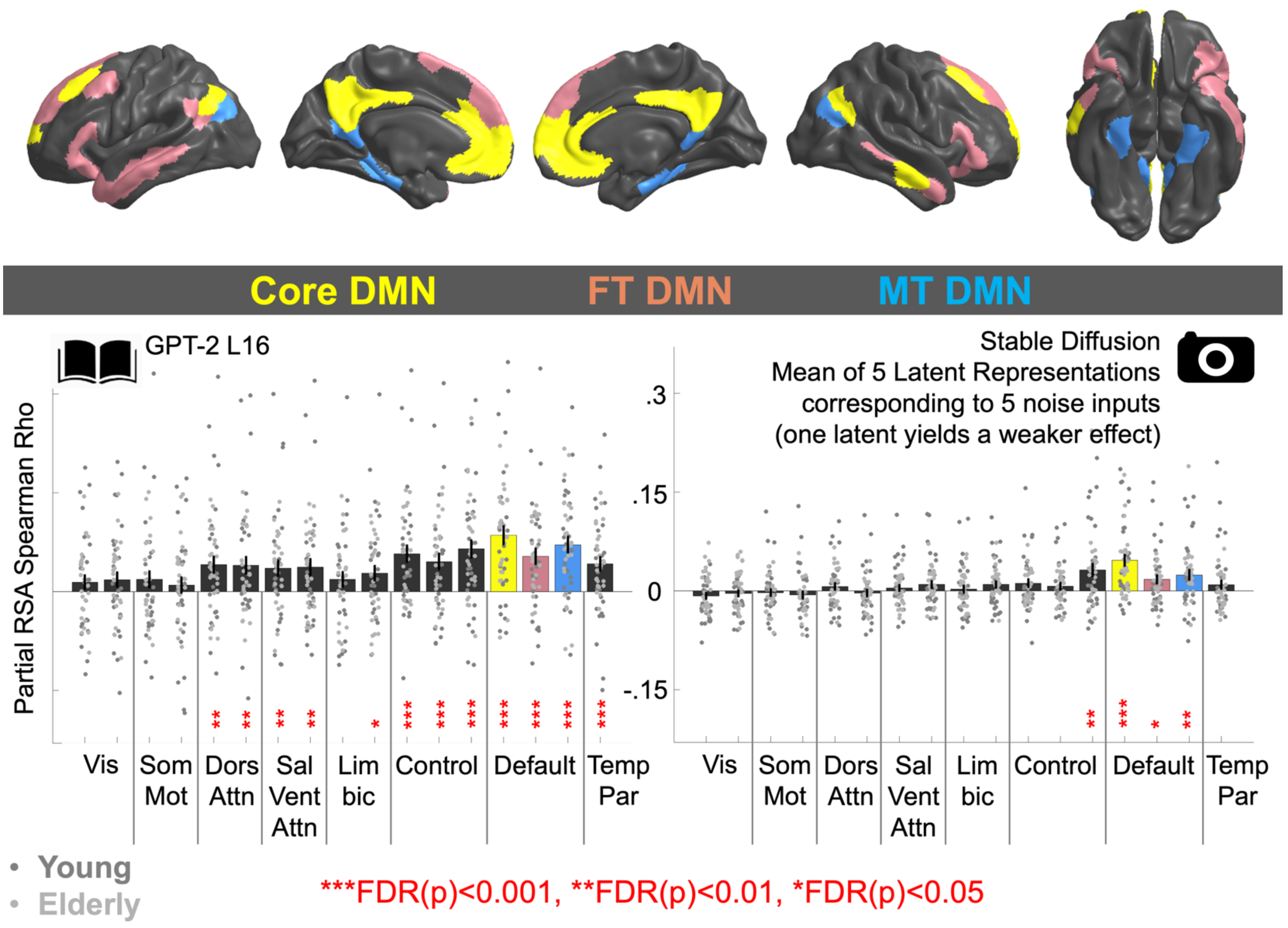


**Supplementary Figure 5.** Repeating **Fig. 2** with Stable Diffusion Latent Representations in place of the Visual model (Stable Diffusion and VGG-16[Fc8]) yields weaker MT-DMN sensitivity

To establish whether MT-DMN activation was captured by the compressed latent representations upon which the Reverse Diffusion process operates (see **Methods**) the analyses of **Fig. 2** were repeated using latent Scenario representations in place of the Visual model. To conduct this analysis, latent representations associated with five synthesized images were vectorized (from a 64*64*4 space), and pointwise averaged to produce a single long vector for each scenario in each participant. In passing, pointwise averaging yielded stronger outcomes than conducting the analysis on a single latent representation per participant – Results not displayed). The Partial RSA analysis was rerun, and partial correlation RSA coefficients corresponding to GPT-2 (Book icon) and the Latent model (Camera Icon) are displayed in the Left and Right plots respectively for fifty participants in each of the seventeen Yeo networks. Partial RSA coefficients in MT-DMN were significantly lower for the Latent representations than for the Visual model (**Fig. 2**), z=2.8, p=0.005 (2-tail), n=50, suggesting that the representational format was a weaker model of brain activation.

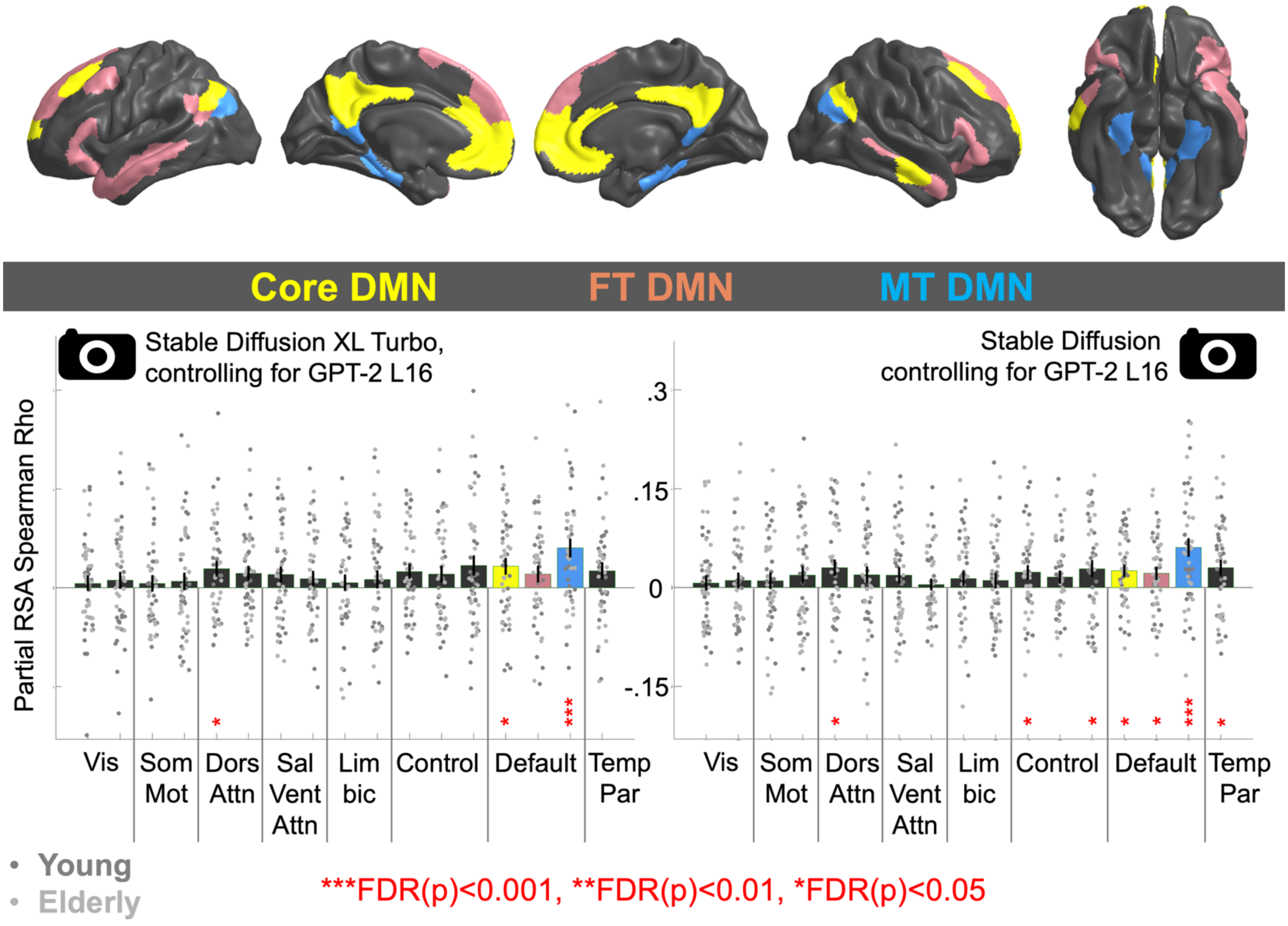


**Supplementary Figure 6. Fig. 2** results are replicated with a different Stable Diffusion model (Stable Diffusion XL Turbo)

Partial correlation RSA coefficients derived from Stable Diffusion XL Turbo (Podell et al. 2023), when controlling for GPT-2) are displayed in the Left plot and the original **Fig. 2** results derived using Stable Diffusion 2.1 (Rombach et al. 2022) are copied into the right plot. Critically, MT-DMN is strongly captured by the Visual model in both cases. Bars indicate mean RSA partial correlation coefficients across participants. Error bars are SEM. P-values correspond to signed-ranks tests computed across all fifty people, adjusted according to False Discovery Rate (Benjamini and Hochberg, 1995).

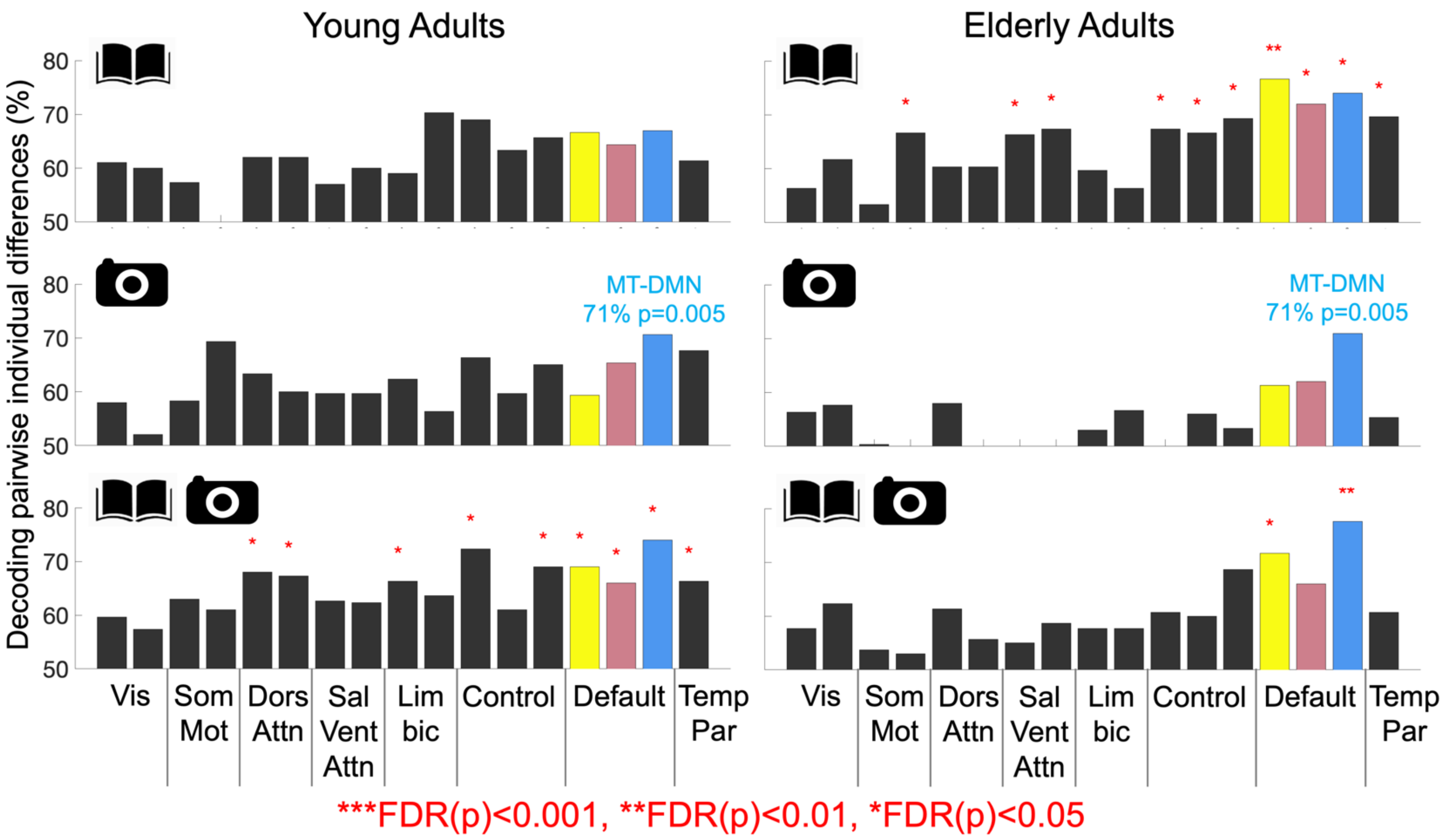


**Supplementary Figure 7.** MT-DMN reflects participant-specific visual content during imagination in young and elderly adults

**A.** Individual-differences associated with the Visual Model were prominent in MT-DMN and present in Core-DMN, whereas GPT-2 captured individual-differences in 11/17 networks, including MT-DMN and Core-DMN.

**B.** Individual-differences in MT-DMN were more accurately discriminated when the Visual Model and GPT-2 were combined, than when GPT-2 was used in isolation.

**Supplementary Table 1.** Experimental sentences and their grammatical structure.

Grammatical elements are tagged in the right column <S> = begin Subject (</S> = end Subject). <V>=Verb. <O>=Direct Object. <IO>=Indirect Object. <Cop>=Copula-phrase. <Adju>=Adjunct.

| | |
|---|---|
| **The family survived the powerful hurricane.** | <S> family </S> <V> survived </V> <O> powerful hurricane </O> |
| **The family was happy.** | <S> family </S> <Cop> happy </Cop> |
| **The family played at the beach.** | <S> family </S> <V> played </V> <Adju> beach </Adju> |
| **The wealthy family celebrated at the party.** | <S> wealthy family </S> <V> celebrated </V> <Adju> party </Adju> |
| **The parent watched the sick child.** | <S> parent </S> <V> watched </V> <O> sick child </O> |
| **The politician visited the family.** | <S> politician </S> <V> visited </V> <O> family </O> |
| **The priest approached the lonely family.** | <S> priest </S> <V> approached </V> <O> lonely family </O> |
| **The parent visited the school.** | <S> parent </S> <V> visited </V> <O> school </O> |
| **The parent shouted at the child.** | <S> parent </S> <V> shouted </V> <IO> child </IO> |
| **The parent bought the magazine.** | <S> parent </S> <V> bought </V> <O> magazine </O> |
| **The happy couple visited the embassy.** | <S> happy couple </S> <V> visited </V> <O> embassy </O> |
| **The couple planned the vacation.** | <S> couple </S> <V> planned </V> <O> vacation </O> |
| **The parent took the cellphone.** | <S> parent </S> <V> took </V> <O> cellphone </O> |
| **The couple laughed at dinner.** | <S> couple </S> <V> laughed </V> <Adju> dinner </Adju> |
| **The couple read on the beach.** | <S> couple </S> <V> read </V> <Adju> beach </Adju> |
| **The wealthy couple left the theater.** | <S> wealthy couple </S> <V> left </V> <O> theater </O> |
| **The child broke the glass in the restaurant.** | <S>child </S> <V> broke </V> <O> glass </O> <Adju> restaurant </Adju> |
| **The happy child found the dime.** | <S>happy child </S> <V> found </V> <O> dime </O> |
| **The child gave the flower to the artist.** | <S> child </S> <V> gave </V> <O> flower </O> <IO> artist </IO> |
| **The child held the soft feather.** | <S> child </S> <V> held </V> <O> soft feather </O> |
| **The angry child threw the book.** | <S> angry child </S> <V> threw </V> <O> book </O> |
| **The girl dropped the shiny dime.** | <S> girl </S> <V> dropped </V> <O> shiny dime </O> |
| **The actor gave the football to the team.** | <S> actor </S> <V> gave </V> <O> football </O> <IO> team </IO> |
| **The commander listened to the soldier.** | <S> commander </S> <V> listened </V> <IO> soldier </IO> |
| **The soldier crossed the field.** | <S> soldier </S> <V> crossed </V> <O> field </O> |
| **The editor drank tea at dinner.** | <S> editor </S> <V> drank </V> <O> tea </O> <Adju> dinner </Adju> |
| **The beach was empty.** | <S> beach </S> <Cop> empty </Cop> |
| **The judge met the mayor.** | <S> judge </S> <V> met </V> <O> mayor </O> |
| **The doctor stole the book.** | <S> doctor </S> <V> stole </V> <O> book </O> |
| **The artist drew the river.** | <S> artist </S> <V> drew </V> <O> river </O> |
| **The window was dusty.** | <S> window </S> <Cop> dusty </Cop> |
| **The teacher worked at the new school.** | <S> teacher </S> <V>worked </V> <Adju> new school </Adju> |
| **The school was famous.** | <S>school </S> <Cop> famous </Cop> |
| **The school was empty during the summer.** | <S> school </S> <Cop> empty </Cop> <Adju> summer </Adju> |
| **The student walked along the long hall.** | <S>student </S> <V> walked </V> <Adju> long hall </Adju> |
| **The young student read at the desk.** | <S>young student </S> <V> read </V> <Adju> desk </Adju> |
| **The small church was near the school.** | <S>small church </S> <Cop> school </Cop> |
| **The teacher used the computer.** | <S>teacher </S> <V> used </V> <O> computer </O> |
| **The army marched past the school.** | <S> army </S> <V> marched </V> <Adju> school </Adju> |
| **The scientist spoke to the student.** | <S>scientist </S> <V> spoke </V> <IO> student </IO> |

| | |
|---|---|
| The engineer gave a book to the student. | <S> engineer </S> <V> gave </V> <O> book </O> <IO> student </IO> |
| The student planned the protest. | <S> student </S> <V> planned </V> <O> protest </O> |
| The teacher broke the small camera. | <S> teacher </S> <V> broke </V> <O> small camera </O> |
| The yellow dog approached the friendly teacher. | <S>yellow dog </S> <V> approached </V> <O> friendly teacher </O> |
| The teacher visited the beach in summer. | <S> teacher </S> <V> visited </V> <O> beach </O> <Adju> summer </Adju> |
| The red pencil was on the desk. | <S>red pencil </S> <Cop> desk </Cop> |
| The team played soccer in spring. | <S>team </S> <V> played </V> <O> soccer </O> <Adju> spring </Adju> |
| The council read the agreement. | <S> council </S> <V> read </V> <O> agreement </O> |
| The mayor dropped the glass. | <S> mayor </S> <V> dropped </V> <O> glass </O> |
| The street was dark. | <S> street </S> <Cop> dark </Cop> |
| The feather was blue. | <S> feather </S> <Cop> blue </Cop> |
| The tree was green. | <S> tree </S> <Cop> green </Cop> |
| The diplomat was wealthy. | <S> diplomat </S> <Cop> wealthy </Cop> |
| The dime was new. | <S> dime </S> <Cop> new </Cop> |
| The girl saw the small bird. | <S> girl </S> <V> saw </V> <O> small bird </O> |
| The small boy feared the storm. | <S> small boy </S> <V> feared </V> <O> storm </O> |
| The mouse ran into the forest. | <S> mouse </S> <V> ran </V> <Adju> forest </Adju> |
| The boat crossed the small lake. | <S> boat </S> <V> crossed </V> <O> small lake </O> |
| The army built the small hospital. | <S> army </S> <V> built </V> <O> small hospital </O> |
| The judge lost the dime. | <S> judge </S> <V> lost </V> <O> dime </O> |
| The man saw the dead mouse. | <S> man </S> <V> saw </V> <O> dead mouse </O> |
| The boy kicked the stone along the street. | <S> boy </S> <V> kicked </V> <O> stone </O> <Adju> street </Adju> |
| The white feather was under the tree. | <S> white feather </S> <Cop> tree </Cop> |
| The dusty feather landed on the highway. | <S> dusty feather </S> <V> landed </V> <Adju> highway </Adju> |
| The cellphone was black. | <S> cellphone </S> <Cop> black </Cop> |
| The fish lived in the river. | <S> fish </S> <V> lived </V> <Adju> river </Adju> |
| The activist dropped the new cellphone. | <S> activist </S> <V> dropped </V> <O> new cellphone </O> |
| The woman bought medicine at the store. | <S> woman </S> <V> bought </V> <O> medicine </O> <Adju> store </Adju> |
| The magazine was yellow. | <S> magazine </S> <Cop> yellow </Cop> |
| The minister found cash at the airport. | <S> minister </S> <V> found </V> <O> cash </O> <Adju> airport </Adju> |
| The businessman laughed in the theater. | <S> businessman </S> <V> laughed </V> <Adju> theater </Adju> |
| The big horse drank from the lake. | <S> big horse </S> <V> drank </V> <Adju> lake </Adju> |
| The pilot was friendly. | <S> pilot </S> <Cop> friendly </Cop> |
| The witness spoke to the lawyer. | <S> witness </S> <V> spoke </V> <IO> lawyer </IO> |
| The minister spoke to the injured patient. | <S> minister </S> <V> spoke </V> <IO> injured patient </IO> |
| The reporter spoke to the loud mob. | <S> reporter </S> <V> spoke </V> <IO> loud mob </IO> |
| The young author spoke to the editor. | <S> young author </S> <V> spoke </V> <IO> editor </IO> |
| The author interviewed the scientist after the flood. | <S> author </S> <V> interviewed </V> <O> scientist </O> <Adju> flood </Adju> |
| The commander negotiated with the council. | <S> commander </S> <V> negotiated </V> <IO> council </IO> |
| The diplomat negotiated at the embassy. | <S> diplomat </S> <V> negotiated </V> <Adju> embassy </Adju> |
| The journalist interviewed the judge. | <S> journalist </S> <V> interviewed </V> <O> judge </O> |

| | |
|---|---|
| **The reporter interviewed the dangerous terrorist.** | <S> reporter </S> <V> interviewed </V> <O> dangerous terrorist </O> |
| **The policeman interviewed the young victim.** | <S> policeman </S> <V> interviewed </V> <O> young victim </O> |
| **The mayor negotiated with the mob.** | <S> mayor </S> <V> negotiated </V> <IO> mob </IO> |
| **The reporter interviewed the politician during the debate.** | <S> reporter </S> <V> interviewed </V> <O> politician </O> <Adju> debate </Adju> |
| **The witness shouted during the trial.** | <S>witness </S> <V> shouted </V> <Adju> trial </Adju> |
| **The artist shouted in the hotel.** | <S> artist </S> <V> shouted </V> <Adju> hotel </Adju> |
| **The diplomat shouted at the soldier.** | <S> diplomat </S> <V> shouted </V> <IO> soldier </IO> |
| **The activist listened to the tired victim.** | <S> activist </S> <V> listened </V> <IO> tired victim </IO> |
| **The mayor listened to the voter.** | <S> mayor </S> <V> listened </V> <IO> voter </IO> |
| **The jury listened to the famous businessman.** | <S> jury </S> <V> listened </V> <IO> famous businessman </IO> |
| **The woman helped the sick tourist.** | <S> woman </S> <V> helped </V> <O> sick tourist </O> |
| **The lonely patient listened to the loud television.** | <S> lonely patient </S> <V> listened </V> <IO> loud television </IO> |
| **The soldier delivered the medicine during the flood.** | <S> soldier </S> <V> delivered </V> <O> medicine </O> <Adju> flood </Adju> |
| **The engineer built the computer.** | <S> engineer </S> <V> built </V> <O> computer </O> |
| **The terrorist stole the car.** | <S> terrorist </S> <V> stole </V> <O> car </O> |
| **The artist found the red ball.** | <S> artist </S> <V> found </V> <O> red ball </O> |
| **The scientist watched the duck.** | <S> scientist </S> <V> watched </V> <O> duck </O> |
| **The flood was dangerous.** | <S> flood </S> <Cop> dangerous </Cop> |
| **The cloud blocked the sun.** | <S> cloud </S> <V> blocked </V> <O> sun </O> |
| **The baseball broke the window.** | <S>baseball </S> <V> broke </V> <O>window </O> |
| **The dog broke the television.** | <S> dog </S> <V> broke </V> <O> television </O> |
| **The angry activist broke the chair.** | <S> angry activist </S> <V> broke </V> <O> chair </O> |
| **The accident destroyed the empty lab.** | <S> accident </S> <V> destroyed </V> <O> empty lab </O> |
| **The accident damaged the yellow car.** | <S> accident </S> <V> damaged </V> <O> yellow car </O> |
| **The hurricane damaged the boat.** | <S> hurricane </S> <V> damaged </V> <O> boat </O> |
| **The storm destroyed the theater.** | <S> storm </S> <V> destroyed </V> <O> theater </O> |
| **The editor damaged the bicycle.** | <S> editor </S> <V> damaged </V> <O> bicycle </O> |
| **The mob damaged the hotel.** | <S> mob </S> <V> damaged </V> <O> hospital </O> |
| **The flood damaged the hospital.** | <S> flood </S> <V> damaged </V> <O> hospital </O> |
| **The horse kicked the fence.** | <S> horse </S> <V> kicked </V> <O> fence </O> |
| **The soldier kicked the door.** | <S> soldier </S> <V> kicked </V> <O> door </O> |
| **The banker was injured in the accident.** | <S> banker </S> <V> injured </V> <Adju> accident </Adju> |
| **The author kicked the desk.** | <S> author </S> <V> kicked </V> <O> desk </O> |
| **The storm was powerful.** | <S> storm </S> <Cop> powerful </Cop> |
| **The doctor helped the injured policeman.** | <S> doctor </S> <V> helped </V> <O> injured policeman </O> |
| **The injured horse slept at night.** | <S> injured horse </S> <V> slept </V> <Adju> night </Adju> |
| **The soldier arrested the injured activist.** | <S>soldier </S> <V> arrested </V> <O> injured activist </O> |
| **The dangerous criminal stole the television.** | <S> dangerous criminal </S> <V> stole </V> <O> television </O> |
| **The doctor bought the used boat.** | <S> doctor </S> <V> bought </V> <O> used boat </O> |
| **The guard opened the window.** | <S> guard </S> <V> opened </V> <O> window </O> |
| **The egg was blue.** | <S> egg </S> <Cop> blue </Cop> |
| **The glass was cold.** | <S> glass </S> <Cop> cold </Cop> |

| | |
|---|---|
| The witness went to the trial. | <S> witness </S> <V> went </V> <Adju> trial </Adju> |
| The trial ended in spring. | <S> trial </S> <V> ended </V> <Adju> spring </Adju> |
| The politician watched the trial. | <S> politician </S> <V> watched </V> <O> trial </O> |
| The reporter wrote about the trial. | <S> reporter </S> <V> wrote </V> <IO> trial </IO> |
| The activist marched at the trial. | <S> activist </S> <V> marched </V> <Adju> trial </Adju> |
| The tired jury left the court. | <S> tired jury </S> <V> left </V> <O> court </O> |
| The jury watched the witness. | <S> jury </S> <V> watched </V> <O> witness </O> |
| The lawyer was friendly. | <S> lawyer </S> <Cop> friendly </Cop> |
| The angry lawyer left the office. | <S>angry lawyer </S> <V> left </V> <O> office </O> |
| The tired lawyer visited the island. | <S> tired lawyer </S> <V> visited </V> <O> island </O> |
| The lawyer drank coffee. | <S> lawyer </S> <V> drank </V> <O> coffee </O> |
| The old judge saw the dark cloud. | <S> old judge </S> <V> saw </V> <O> dark cloud </O> |
| The judge stayed at the hotel during the vacation. | <S> judge </S> <V> stayed </V> <IO> hotel </IO> <Adju> vacation </Adju> |
| The policeman arrested the angry driver. | <S> policeman </S> <V> arrested </V> <O> angry driver </O> |
| The tired patient slept in the dark hospital. | <S> tired patient </S> <V> slept </V> <Adju> dark hospital </Adju> |
| The man read the newspaper in church. | <S> man </S> <V> read </V> <O> newspaper </O> <Adju> church </Adju> |
| The criminal wanted cash. | <S> criminal </S> <V> wanted </V> <O> cash </O> |
| The clever scientist worked at the lab. | <S> clever scientist </S> <V> worked </V> <Adju> lab </Adju> |
| The editor gave cash to the driver. | <S> editor </S> <V> gave </V> <O> cash </O> <IO> driver </IO> |
| The green car crossed the bridge. | <S> green car </S> <V> crossed </V> <O> bridge </O> |
| The vacation was peaceful. | <S> vacation </S> <Cop> peaceful </Cop> |
| The duck lived at the lake. | <S> duck </S> <V> lived </V> <IO> lake </IO> |
| The bird landed on the bridge. | <S> bird </S> <V> landed </V> <Adju> bridge </Adju> |
| The protest was loud. | <S> protest </S> <Cop> loud </Cop> |
| The voter went to the protest. | <S> voter </S> <V> went </V> <IO> protest </IO> |
| The council feared the protest. | <S> council </S> <V> feared </V> <O> protest </O> |
| The banker watched the peaceful protest. | <S> banker </S> <V> watched </V> <O> peaceful protest </O> |
| The mob approached the embassy. | <S> mob </S> <V> approached </V> <O> embassy </O> |
| The mob was dangerous. | <S> mob </S> <Cop> dangerous </Cop> |
| The reporter met the angry doctor. | <S> reporter </S> <V> met </V> <O> angry doctor </O> |
| The voter read about the election. | <S> voter </S> <V> read </V> <IO> election </IO> |
| The politician celebrated at the hotel. | <S> politician </S> <V> celebrated </V> <Adju> hotel </Adju> |
| The wealthy politician liked coffee. | <S> wealthy politician </S> <V> liked </V> <O> coffee </O> |
| The worker fixed the door at the church. | <S> worker </S> <V> fixed </V> <O> door </O> <Adju> church </Adju> |
| The corn grew in spring. | <S> corn </S> <V> grew </V> <Adju> spring </Adju> |
| The victim feared the criminal. | <S>victim </S> <V> feared </V> <O> criminal </O> |
| The young engineer worked in the office. | <S> young engineer </S> <V> worked </V> <Adju> office </Adju> |
| The tourist was friendly. | <S> tourist </S> <Cop> friendly </Cop> |
| The baseball was in the office. | <S> baseball </S> <Cop> office </Cop> |
| The used book was on the table. | <S> used book </S> <Cop> table </Cop> |
| The magazine was in the car. | <S> magazine </S> <Cop> car </Cop> |
| The bridge survived the flood. | <S> bridge </S> <V> survived </V> <O> flood </O> |
| The old doctor walked through the hospital. | <S> old doctor </S> <V> walked </V> <Adju> hospital </Adju> |

| | |
|---|---|
| The patient survived. | <S> patient </S> <V>survived </V> |
| The patient put the medicine in the cabinet. | <S> patient </S> <V> put </V> <O> medicine </O> <IO> cabinet </IO> |
| The medicine was on the table. | <S>medicine </S> <Cop> table </Cop> |
| The famous diplomat left the hospital. | <S> famous diplomat </S> <V> left </V> <O> hospital </O> |
| The commander opened the heavy door. | <S> commander </S> <V> opened </V> <O> heavy door </O> |
| The banker bought the expensive boat. | <S>banker </S> <V> bought </V> <O> expensive boat </O> |
| The storm ended during the morning. | <S>storm </S> <V> ended </V> <Adju> morning </Adju> |
| The car approached the river. | <S> car </S> <V> approached </V> <O> river </O> |
| The door was blue. | <S>door </S> <Cop> blue </Cop> |
| The farmer liked soccer. | <S> farmer </S> <V> liked </V> <O> soccer </O> |
| The engineer walked in the peaceful park. | <S> engineer </S> <V> walked </V> <Adju> peaceful park </Adju> |
| The horse walked through the green field. | <S> horse </S> <V> walked </V> <Adju> green field </Adju> |
| The wealthy author walked into the office. | <S> wealthy author </S> <V> walked </V> <Adju> office </Adju> |
| The young policeman walked to the theater. | <S> young policeman </S> <V> walked </V> <Adju> theater </Adju> |
| The artist hiked along the mountain. | <S> artist </S> <V> hiked </V> <Adju> mountain </Adju> |
| The tourist hiked through the forest. | <S>tourist </S> <V> hiked </V> <Adju> forest </Adju> |
| The editor carried the magazine to the meeting. | <S> editor </S> <V> carried </V> <O> magazine </O> <IO> meeting </IO> |
| The dog ran in the park. | <S> dog </S> <V> ran </V> <Adju> park </Adju> |
| The woman took the flower from the field. | <S> woman </S> <V> took </V> <O> flower </O> <Adju> field </Adju> |
| The street was empty at night. | <S> street </S> <Cop> empty </Cop> <Adju> night </Adju> |
| The wealthy farmer fed the horse. | <S> wealthy farmer </S> <V> fed </V> <O>horse </O> |
| The diplomat bought the aggressive dog. | <S> diplomat </S> <V> bought </V> <O> aggressive dog </O> |
| The dog drank water. | <S> dog </S> <V> drank </V> <O> water </O> |
| The tree grew in the park. | <S> tree </S> <V> grew </V> <Adju> park </Adju> |
| The commander ate chicken at dinner. | <S> commander </S> <V> ate </V> <O> chicken </O> <Adju> dinner </Adju> |
| The dog ate the egg. | <S> dog </S> <V> ate </V> <O> egg </O> |
| The computer was new. | <S> computer </S> <Cop> new </Cop> |
| The company delivered the computer. | <S> company </S> <V> delivered </V> <O> computer </O> |
| The computer was on the desk. | <S> computer </S> <Cop> desk </Cop> |
| The businessman lost the computer at the airport. | <S> businessman </S> <V> lost </V> <O> computer </O> <Adju> airport </Adju> |
| The expensive camera was in the lab. | <S> expensive camera </S> <Cop> lab </Cop> |
| The reporter ate at the new restaurant. | <S> reporter </S> <V> ate </V> <Adju> new restaurant </Adju> |
| The minister lost the spiritual magazine. | <S> minister </S> <V> lost </V> <O> spiritual magazine </O> |
| The boy held the football. | <S> boy </S> <V> held </V> <O> football </O> |
| The bird was red. | <S> bird </S> <Cop> red </Cop> |
| The cloud was white. | <S> cloud </S> <Cop> white </Cop> |
| The yellow bird flew over the field. | <S> yellow bird </S> <V> flew </V> <Adju> field </Adju> |
| The flower was yellow. | <S> flower </S> <Cop> yellow </Cop> |
| The green duck slept under the tree. | <S> green duck </S> <V> slept </V> <Adju> tree </Adju> |
| The girl saw a horse in the park. | <S> girl </S> <V> saw </V> <O> horse </O> <Adju> park </Adju> |
| The duck flew. | <S> duck </S> <V> flew </V> |
| The man lost the ticket to soccer. | <S> man </S> <V> lost </V> <O> soccer ticket </O> |
| The team celebrated. | <S> team </S> <V> celebrated </V> |

| | |
|---|---|
| **The red plane flew through the cloud.** | <S> red plane </S> <V> flew </V> <Adju> cloud </Adju> |
| **The summer was hot.** | <S> summer </S> <Cop> hot </Cop> |
| **The bicycle blocked the green door.** | <S> bicycle </S> <V> blocked </V> <O> green door </O> |
| **The park was empty in winter.** | <S> park </S> <Cop> empty </Cop> <Adju> winter </Adju> |
| **The driver wanted cold tea.** | <S> driver </S> <V> wanted </V> <O> cold tea </O> |
| **The minister visited the prison.** | <S> minister </S> <V> visited </V> <O>prison </O> |
| **The tourist ate bread on vacation.** | <S> tourist </S> <V> ate </V> <O> bread </O> <Adju> vacation </Adju> |
| **The tourist went to the restaurant.** | <S> tourist </S> <V> went </V> <IO> restaurant </IO> |
| **The tourist found a bird in the theater.** | <S> tourist </S> <V> found </V> <O> bird </O> <Adju> theater </Adju> |
| **The old farmer ate at the expensive hotel.** | <S> old farmer </S> <V> ate </V> <Adju> expensive hotel </Adju> |
| **The aggressive team took the baseball.** | <S> aggressive team </S> <V> took </V> <O> baseball </O> |
| **The duck was aggressive.** | <S> duck </S> <Cop> aggressive </Cop> |
| **The chicken was expensive at the restaurant.** | <S> chicken </S> <Cop> expensive </Cop> <Adju>restaurant </Adju> |
| **The artist liked chicken.** | <S> artist </S> <V> liked </V> <O> chicken </O> |
| **The restaurant was loud at night.** | <S> restaurant </S> <Cop> loud </Cop> <Adju>night </Adju> |
| **The woman left the restaurant after the storm.** | <S> woman </S> <V> left </V> <O>restaurant </O> <Adju> storm </Adju> |
| **The banker drank cold water.** | <S> banker </S> <V> drank </V> <O> cold water </O> |
| **The coffee was hot.** | <S> coffee </S> <Cop> hot </Cop> |
| **The boy threw the baseball over the fence.** | <S> boy </S> <V> threw </V> <O> baseball </O> <Adju> fence </Adju> |
| **The policeman read the newspaper.** | <S> policeman </S> <V> read </V> <O> newspaper </O> |
| **The criminal put the book on the desk.** | <S> criminal </S> <V> put </V> <O> book </O> <IO> desk </IO> |
| **The man saw the fish in the river.** | <S> man </S> <V> saw </V> <O> fish </O> <Adju> river </Adju> |
| **The happy girl played in the forest.** | <S> happy girl </S> <V> played </V> <Adju> forest </Adju> |
| **The young girl played soccer.** | <S> young girl </S> <V> played </V> <O> soccer </O> |
| **The old man threw the stone into the lake.** | <S> old man </S> <V> threw </V> <O> stone </O> <Adju> lake </Adju> |
| **The team lost the football in the forest.** | <S> team </S> <V> lost </V> <O> football </O> <Adju> forest </Adju> |
| **The businessman slept on the expensive bed.** | <S> businessman </S> <V> slept </V> <Adju> expensive bed </Adju> |
| **The businessman watched soccer.** | <S> businessman </S> <V> watched </V> <O> soccer </O> |
| **The guard slept near the door.** | <S> guard </S> <V> slept </V> <Adju> door </Adju> |
| **The artist kicked the football.** | <S> artist </S> <V> kicked </V> <O> football </O> |
| **The ticket was on the red desk.** | <S> ticket </S> <Cop> red desk </Cop> |

### fMRIPrep Boilerplate Template

### Copyright Waiver

The below boilerplate text was automatically generated by fMRIPrep[10] with the express intention that users should copy and paste this text into their manuscripts unchanged. It is released under the CC0 license.

#### Boilerplate

Results included in this manuscript come from preprocessing performed using fMRIPrep 20.2.1 (Esteban, Markiewicz, et al. (2018)[10]; Esteban, Blair, et al. (2018)[11]; RRID:SCR_016216), which is based on Nipype 1.5.1 (Gorgolewski et al. (2011)[12]; Gorgolewski et al. (2018)[13]; RRID:SCR_002502).

#### Anatomical data preprocessing

A total of 1 T1-weighted (T1w) images were found within the input BIDS dataset.The T1-weighted (T1w) image was corrected for intensity non-uniformity (INU) with N4BiasFieldCorrection (Tustison et al. 2010[14]), distributed with ANTs 2.3.3 (Avants et al. 2008[15], RRID:SCR_004757), and used as T1w-reference throughout the workflow. The T1w-reference was then skull-stripped with a Nipype implementation of the antsBrainExtraction.sh workflow (from ANTs), using OASIS30ANTs as target template. Brain tissue segmentation of cerebrospinal fluid (CSF), white-matter (WM) and gray-matter (GM) was performed on the brain-extracted T1w using fast (FSL 5.0.9, RRID:SCR_002823, Zhang, Brady, and Smith 2001[16]). Brain surfaces were reconstructed using recon-all (FreeSurfer 6.0.1, RRID:SCR_001847, Dale, Fischl, and Sereno 1999[17]), and the brain mask estimated previously was refined with a custom variation of the method to reconcile ANTs-derived and FreeSurfer-derived segmentations of the cortical gray-matter of Mindboggle (RRID:SCR_002438, Klein et al. 2017[18]). Volume-based spatial normalization to one standard space (MNI152NLin2009cAsym) was performed through nonlinear registration with antsRegistration (ANTs 2.3.3), using brain-extracted versions of both T1w reference and the T1w template. The following template was selected for spatial normalization: ICBM 152 Nonlinear Asymmetrical template version 2009c [Fonov et al. (2009)[19], RRID:SCR_008796; TemplateFlow ID: MNI152NLin2009cAsym],

#### Functional data preprocessing

For each of the 1 BOLD runs found per subject (across all tasks and sessions), the following preprocessing was performed. First, a reference volume and its skull-stripped version were generated using a custom methodology of fMRIPrep. Susceptibility distortion correction (SDC) was omitted. The BOLD reference was then co-registered to the T1w reference using bbregister (FreeSurfer) which implements boundary-based registration (Greve and Fischl 2009[20]). Co-registration was configured with six degrees of freedom. Head-motion parameters with respect to the BOLD reference (transformation matrices, and six corresponding rotation and translation parameters) are estimated before any spatiotemporal filtering using mcflirt (FSL 5.0.9, Jenkinson et al. 2002[21]). BOLD runs were slice-time corrected using 3dTshift from AFNI 20160207 (Cox and Hyde 1997[22], RRID:SCR_005927). The BOLD time-series (including slice-timing correction when applied) were resampled onto their original, native space by applying the transforms to correct for head-motion. These resampled BOLD time-series will be referred to as preprocessed BOLD in original space, or just preprocessed BOLD. The BOLD time-series were resampled into standard space, generating a preprocessed BOLD run in MNI152NLin2009cAsym space. First, a reference volume and its skull-stripped version were generated using a custom methodology of fMRIPrep. Several confounding time-series were calculated based on the preprocessed BOLD: framewise displacement (FD), DVARS and three region-wise global signals. FD was computed using two formulations following Power (absolute sum of relative motions, Power et al. (2014)[23]) and Jenkinson (relative root mean square displacement between affines, Jenkinson et al. (2002)[21]). FD and DVARS are calculated for each functional run, both using their implementations in Nipype (following the definitions by Power et al. 2014[23]). The three global signals are extracted within the CSF, the WM, and the whole-brain masks. Additionally, a set of physiological regressors were extracted to allow for component-based noise correction (CompCor, Behzadi et al. 2007[24]). Principal components are estimated after high-pass filtering the preprocessed BOLD time-series (using a discrete cosine filter with 128s cut-off) for the two CompCor variants: temporal (tCompCor) and anatomical (aCompCor). tCompCor components are then calculated from the top 2% variable voxels within the brain mask. For aCompCor, three probabilistic masks (CSF, WM and combined CSF+WM) are generated in anatomical space. The implementation differs from that of Behzadi et al. in that instead of eroding the masks by 2 pixels on BOLD space, the aCompCor masks are subtracted a mask of pixels that likely contain a volume fraction of GM. This mask is obtained by dilating a GM mask extracted from the FreeSurfer's aseg segmentation, and it ensures components are not extracted from voxels containing a minimal fraction of GM. Finally, these masks are resampled into BOLD space and binarized by thresholding at 0.99 (as in the original implementation). Components are also calculated separately within the WM and CSF masks. For each CompCor decomposition, the k components with the largest singular values are retained, such that the retained components' time series are sufficient to explain 50 percent of variance across the nuisance mask (CSF, WM, combined, or temporal). The remaining components are dropped from consideration. The head-motion estimates calculated in the correction step were also placed within the corresponding confounds file. The confound time series derived from head motion estimates and global signals were expanded with the inclusion of temporal derivatives and quadratic terms for each (Satterthwaite et al. 2013[25]). Frames that exceeded a threshold of 0.5 mm FD or 1.5 standardised DVARS were annotated as motion outliers. All resamplings can be performed with a single interpolation step by composing all the pertinent transformations (i.e. head-motion transform matrices, susceptibility distortion correction when available, and co-registrations to anatomical and output spaces). Gridded (volumetric) resamplings were performed using antsApplyTransforms (ANTs), configured with Lanczos interpolation to minimize the smoothing effects of other kernels (Lanczos 1964[26]). Non-gridded (surface) resamplings were performed using mri_vol2surf (FreeSurfer).

Many internal operations of fMRIPrep use Nilearn 0.6.2 (Abraham et al. 2014[27], RRID:SCR_001362), mostly within the functional processing workflow. For more details of the pipeline, see the section corresponding to workflows in fMRIPrep's documentation.

### References


1. Benjamini Y, Hochberg Y. 1995. Controlling the false discovery rate: a practical and powerful approach to multiple testing. Journal of the Royal statistical society: series B (Methodological). 57(1):289-300.
2. Mitchell TM, Shinkareva SV, Carlson A, Chang K-M, Malave VL, Mason RA, Just MA. 2008. Predicting human brain activity associated with the meaning of nouns. Science. 320:1191–1195.
3. Kriegeskorte N, Mur M, Bandettini PA. 2008. Representational similarity analysis-connecting the branches of systems neuroscience. Frontiers in systems neuroscience. 24;2:249.
4. Yeo BT, Krienen FM, Sepulcre J, Sabuncu MR, Lashkari D, Hollinshead M, Roffman JL, Smoller JW, Zöllei L, Polimeni JR, Fischl B, Hesheng L, Buckner RL. 2011. The organization of the human cerebral cortex estimated by intrinsic functional connectivity. Journal of Neurophysiology. 106(3):1125.
5. Schaefer A, Kong R, Gordon EM, Laumann TO, Zuo XN, Holmes AJ, Eickhoff SB, Yeo BTT. 2018. Local-Global parcellation of the human cerebral cortex from intrinsic functional connectivity MRI. Cerebral Cortex, 29:3095-3114.
6. Yarkoni T, Poldrack RA, Nichols TE, Van Essen DC, Wager TD. 2011. Large-scale automated synthesis of human functional neuroimaging data. Nature Methods. 8(8):665-70.
7. Vos de Wael R, Benkarim O, Paquola C, Lariviere S, Royer J, Tavakol S, Xu T, Hong SJ, Langs G, Valk S, Misic B, Milham M, Margulies D, Smallwood J, Bernhardt BC. 2020. BrainSpace: a toolbox for the analysis of macroscale gradients in neuroimaging and connectomics datasets. Communications Biology. 3:103. doi: 10.1038/s42003-020-0794-7.
8. Anderson AJ, Kiela D, Binder JR, Fernandino L, Humphries CJ, Conant LL, Raizada RDS, Grimm S, Lalor EC. 2021. Deep artificial neural networks reveal a distributed cortical network encoding propositional sentence-level meaning. Journal of Neuroscience. 41(18):4100-19.
9. Radford A, Wu J, Child R, Luan D, Amodei D, Sutskever I. 2019. Language models are unsupervised multitask learners. OpenAI blog, 1(8):9.
10. Esteban O, Markiewicz CJ, Blair RW, Moodie CA, Isik AI, Erramuzpe A, Kent JD, Goncalves M, DuPre E, Snyder M, Oya H, Ghosh SS, Wright J, Durnez J, Poldrack RA, Gorgolewski KJ. fMRIPrep: a robust preprocessing pipeline for functional MRI. Nat Meth. 2018; doi:10.1038/s41592-018-0235-4
11. Esteban O, Blair RW, Markiewicz CJ, Berleant SL, Moodie C, Ma F, Isik AI, et al. 2018. FMRIPrep. Software. Zenodo. https://doi.org/10.5281/zenodo.852659.10.Gorgolewski, K., C. D. Burns, C. Madison, D. Clark, Y. O. Halchenko, M. L. Waskom, and S. Ghosh. 2011. Nipype: A Flexible, Lightweight and Extensible Neuroimaging Data Processing Framework in Python. Frontiers in Neuroinformatics 5: 13. https://doi.org/10.3389/fninf.2011.00013.
12. Gorgolewski K, Burns CD, Madison C, Clark D, Halchenko YO, Waskom ML, Ghosh SS. 2011. Nipype: a flexible, lightweight and extensible neuroimaging data processing framework in python. Frontiers in neuroinformatics. ;5:13.
13. Gorgolewski KJ, Esteban O, Ellis DG, Notter MP, Ziegler E, Johnson H, Hamalainen C, Yvernault B, Burns C, Manhães-Savio A, Jarecka D, Markiewicz CJ, Salo T, Clark D, Waskom M, Wong J, Modat M, Dewey BE, Clark MG, Dayan M, Loney F, Madison C, Gramfort A, Keshavan A, Berleant S, Pinsard B, Goncalves M, Clark D, Cipollini B, Varoquaux G, Wassermann D, Rokem A, Halchenko YO, Forbes J, Moloney B, Malone IB, Hanke M, Mordom D, Buchanan C, Pauli WM, Huntenburg JM, Horea C, Schwartz Y, Tungaraza R, Iqbal S, Kleesiek J, Sikka S, Frohlich C, Kent J, Perez-Guevara M, Watanabe A, Welch D, Cumba C, Ginsburg D, Eshaghi A, Kastman E, Bougacha S, Blair R, Acland B, Gillman A, Schaefer A, Nichols BN, Giavasis S, Erickson D, Correa C, Ghayoor A, Küttner R, Haselgrove C, Zhou D, Craddock RC, Haehn D, Lampe L, Millman J, Lai J, Renfro M, Liu S, Stadler J, Glatard T, Kahn AE, Kong X-Z, Triplett W, Park A, McDermottroe C, Hallquist M, Poldrack R, Perkins LN, Noel M, Gerhard S, Salvatore J, Mertz F, Broderick

W, Inati S, Hinds O, Brett M, Durnez J, Tambini A, Rothmei S, Andberg SK, Cooper G, Marina A, Mattfeld A, Urchs S, Sharp P, Matsubara K, Geisler D, Cheung B, Floren A, Nickson T, Pannetier N, Weinstein A, Dubois M, Arias J, Tarbert C, Schlamp K, Jordan K, Liem F, Saase V, Harms R, Khanuja R, Podranski K, Flandin G, Papadopoulos Orfanos D, Schwabacher I, McNamee D, Falkiewicz M, Pellman J, Linkersdörfer J, Varada J, Pérez-García F, Davison A, Shachnev D, Ghosh S. 2018. Nipype. Software. Zenodo. https://doi.org/10.5281/zenodo.596855.

14. Tustison NJ, Avants BB, Cook PA, Zheng Y, Egan A, Yushkevich PA, Gee JC. N4ITK: improved N3 bias correction. 2010. IEEE Trans Med Imaging. 29(6):1310–20. doi:10.1109/TMI.2010.2046908.
15. Avants BB, Epstein CL, Grossman M, Gee JC. 2008. Symmetric diffeomorphic image registration with cross-correlation: evaluating automated labeling of elderly and neurodegenerative brain. Med Image Anal. 2008 12(1):26–41. doi:10.1016/j.media.2007.06.004.
16. Zhang Y, Brady M, Smith S. Segmentation of brain MR images through a hidden Markov random field model and the expectation-maximization algorithm. 2001. IEEE Trans Med Imaging. 20(1):45–57. doi:10.1109/42.906424.
17. Dale A, Fischl B, Sereno MI. 1999. Cortical Surface-Based Analysis: I. Segmentation and Surface Reconstruction. Neuroimage. 9(2):179–94. doi:10.1006/nimg.1998.0395.
18. Klein A, Ghosh SS, Bao FS, Giard J, Häme Y, Stavsky E, Lee N, et al. 2017. Mindboggling Morphometry of Human Brains. PLOS Computational Biology 13 (2): e1005350. https://doi.org/10.1371/journal.pcbi.1005350.
19. Fonov, VS, AC Evans, RC McKinstry, CR Almli, and DL Collins. 2009. Unbiased Nonlinear Average Age-Appropriate Brain Templates from Birth to Adulthood. NeuroImage 47, Supplement 1: S102. https://doi.org/10.1016/S1053-8119(09)70884-5.
20. Greve, Douglas N, and Bruce Fischl. 2009. Accurate and Robust Brain Image Alignment Using Boundary-Based Registration. NeuroImage 48 (1): 63–72. https://doi.org/10.1016/j.neuroimage.2009.06.060.
21. Jenkinson M, Bannister P, Brady M, Smith S. 2002. Improved optimization for the robust and accurate linear registration and motion correction of brain images. Neuroimage. 17(2):825–41. doi:10.1006/nimg.2002.1132.
22. Cox RW, Hyde JS. 1997. Software Tools for Analysis and Visualization of fMRI Data. NMR in Biomedicine 10 (4-5): 171–78. https://doi.org/10.1002/(SICI)1099-1492(199706/08)10:4/5<171::AID-NBM453>3.0.CO;2-L.
23. Power JD, Mitra A, Laumann TO, Snyder AZ, Schlaggar BL, Petersen SE. 2013. Methods to detect, characterize, and remove motion artifact in resting state fMRI. Neuroimage. ;84:320–41. doi:10.1016/j.neuroimage.2013.08.048.
24. Behzadi Y, Restom K, Liau J, Liu TT. 2007. A component based noise correction method (CompCor) for BOLD and perfusion based fMRI. Neuroimage. 37(1):90–101. doi:10.1016/j.neuroimage.2007.04.042.
25. Satterthwaite TD, Elliott MA, Gerraty RT, Ruparel K, Loughead J, Calkins ME, Eickhoff SB, Hakonarson H, Gur RC, Gur RE, Wolf DH. An improved framework for confound regression and filtering for control of motion artifact in the preprocessing of resting-state functional connectivity data. 2013. NeuroImage 64 (1): 240–56. https://doi.org/10.1016/j.neuroimage.2012.08.052.
26. Lanczos C. 1964. Evaluation of Noisy Data. Journal of the Society for Industrial and Applied Mathematics Series B Numerical Analysis 1 (1): 76–85. https://doi.org/10.1137/0701007.
27. Abraham A, Pedregosa F, Eickenberg M, Gervais P, Mueller A, Kossaifi J, Gramfort A, Thirion B, Varoquaux G. Machine learning for neuroimaging with scikit-learn. Front in Neuroinf 8:14. 2014. doi:10.3389/fninf.2014.00014.
